# Intestine-enriched *apolipoprotein b* orthologs are required for stem cell differentiation and regeneration in planarians

**DOI:** 10.1101/2021.03.04.433904

**Authors:** Lily L. Wong, Christina G. Bruxvoort, Nicholas I. Cejda, Jannette Rodriguez Otero, David J. Forsthoefel

**Affiliations:** Genes and Human Disease Research Program, Oklahoma Medical Research Foundation, Oklahoma City, Oklahoma; Current address: Arthritis and Clinical Immunology Research Program, Oklahoma Medical Research Foundation; Department of Pathology, University of Oklahoma Health Sciences Center; and Department of Veteran Affairs Medical Center - Research Services, Oklahoma City, Oklahoma; Howard Hughes Medical Institute, Department of Cell and Developmental Biology, University of Illinois at Urbana-Champaign, Urbana, Illinois; Department of Education, Universidad Interamericana de Puerto Rico, San Juan, Puerto Rico; Department of Cell Biology, University of Oklahoma Health Sciences Center, Oklahoma City, Oklahoma

**Author notes:** Correspondence: David J. Forsthoefel, Ph.D. Authors contributed equally (Lily L. Wong, Christina G. Bruxvoort).

## Abstract

Lipid metabolism plays an instructive role in regulating stem cell state and differentiation. However, the roles of lipid mobilization and utilization in stem cell-driven regeneration are unclear. Planarian flatworms readily restore missing tissue due to injury-induced activation of pluripotent somatic stem cells called neoblasts. Here, we identify two intestine-enriched orthologs of *apolipoprotein b, apob-1* and *apob-2,* which mediate transport of neutral lipid stores from the intestine to target tissues including neoblasts, and are required for tissue homeostasis and regeneration. Inhibition of *apob* function by RNAi causes head regression and lysis in uninjured animals, and delays body axis re-establishment and regeneration of multiple organs in amputated fragments. Furthermore, *apob* RNAi causes expansion of the population of differentiating neoblast progeny and dysregulates expression of genes enriched in differentiating and mature cells in eight major cell type lineages. We conclude that intestine-derived lipids serve as a source of metabolites required for neoblast differentiation.

## Introduction

Regeneration requires metabolites and energy for cell proliferation, differentiation, migration, and growth. Currently, specific mechanisms by which various metabolites are produced, transported, and utilized to promote regeneration are not well understood. In animals, neutral lipids (NLs) (e.g., triglycerides and cholesteryl esters) are stored in intracellular lipid droplets, which serve as central organelles for energy and lipid homeostasis^1, 2^. Lipid stores can be mobilized either by NL packaging and export in lipoprotein particles (LPs), or by lipolysis and secretion as fatty acids (FAs)^3–5^. Mobilization enables transport to other tissues, for example between intestine, liver, and peripheral tissues such as brain and muscle in vertebrates, or between the fat body, intestine, nervous system, and imaginal discs in *Drosophila*^6–8^. Upon lipase-mediated hydrolysis of NLs, the FAs and cholesterol produced serve as building blocks for membrane biosynthesis, substrates for energy via fatty acid beta oxidation, sources of acetyl-CoA and other precursors for chromatin remodeling, and precursors of signaling molecules like eicosanoids and sphingolipids^9–11^.

Considerable evidence indicates that disruption of lipid catabolism, biosynthesis, and transport can dysregulate stem cell pluripotency, proliferation, and differentiation^12–14^. For example, inhibiting fatty acid oxidation (FAO, the process of converting fatty acids to acetyl-CoA) causes symmetric production of differentiating progeny and stem cell depletion in the mammalian hematopoietic lineage^15^. By contrast, in the adult mouse hippocampus, inhibiting FAO promotes exit from quiescence and proliferation of neural stem/progenitor cells in the adult mouse hippocampus^16^. In *Drosophila*, compromising lipolysis and/or FAO causes intestinal stem cell necrosis^17^, and loss of germline stem cells^18^. On the other hand, FAO-mediated production of acetyl-CoA (a precursor for histone acetylation) is required for differentiation, but not maintenance, of quiescent hematopoietic stem cells in *Drosophila* larvae, illustrating the growing recognition of the intersection of lipid metabolism and epigenetic regulation^19^. Perturbing lipid synthesis and delivery also dysregulates stem cell state and differentiation dynamics. For example, inhibiting fatty acid synthase (required for *de novo* lipogenesis) reduces proliferation by adult mouse neural stem and progenitor cells^20^. Similarly, mutations in *hydroxysteroid (17-beta) dehydrogenase 7*, a regulator of cholesterol biosynthesis, cause premature differentiation of neural progenitors during development^21^. In culture, depriving human pluripotent stem cells of extrinsic lipids promotes a “naive-to-primed” intermediate state, demonstrating the importance of exogenous lipid availability for regulation of stem cell state^22^.

Because stem and progenitor cell activation, proliferation, and differentiation are central to regeneration, these diverse observations underline the potential importance of lipid metabolism during regenerative growth. Currently, however, evidence that lipid transport and utilization influence regeneration is limited. In zebrafish, *leptin b* (a hormonal regulator of systemic lipid metabolism) is one of the most upregulated genes during fin and heart regeneration^23^, and *apolipoprotein E* (a regulator of NL transport in LPs) is upregulated during fin regeneration^24^. In the axolotl *Ambystoma mexicanum*, genes involved in steroid, cholesterol, and fatty acid metabolism are among the most upregulated at later stages of limb regeneration, when differentiation of cartilage and muscle progenitors occurs^25^. In a primary cell culture model of nerve injury, rat retinal ganglion cells regrow projections after axotomy more efficiently in the presence of glial-derived LPs, reinforcing the role of cholesterol in axon regeneration^26^. In mice, *low density lipoprotein receptor* deficiency delays liver regeneration and reduces hepatocyte proliferation^27^. Similarly, elevated FA levels (induced by lipoprotein lipase overexpression) causes lipotoxicity and compromises skeletal muscle regeneration^28^, while inhibition of peroxisomal FAO induces differentiation of myogenic satellite cells and muscle hypertrophy during regeneration^29^. During skin wound repair, inhibiting triglyceride lipase-mediated lipolysis by dermal adipocytes compromises recruitment of inflammatory macrophages and adipocyte fate-switching to extracellular-matrix-secreting myofibroblasts^30^.

Although these intriguing observations point to the importance of lipid metabolism, there is scant functional understanding of how lipid transport and utilization influence stem cell regulation during regeneration, particularly in emerging animal models with extensive regenerative capacity. Planarians are freshwater flatworms capable of whole-body regeneration, an ability conferred by pluripotent somatic stem cells called neoblasts that divide and differentiate to replace damaged and lost tissues after amputation^31, 32^. Diet-derived NLs are stored in the planarian intestine in lipid droplets, suggesting the intestine is a major lipid storage organ, as in *Drosophila* and *C. elegans*^6, 33–35^. Planarian intestinal lipids have been proposed to be a source of metabolites during extended fasting, as well as regeneration^34, 36, 37^. However, mechanisms by which lipid secretion is controlled, and whether delivery to neoblasts or their progeny is functionally required for regeneration, have not been investigated.

In this study, we show that two intestine-enriched *apolipoprotein b* (*apob*) orthologs are required for NL transport from the intestine to neoblasts and their progeny, and that ApoB function is required for stem cell differentiation and regeneration in the planarian *Schmidtea mediterranea*. In mammals and insects, ApoB and ApoB-like proteins mediate trafficking of NLs in LPs from digestive and storage organs to peripheral target tissues^5, 38^. Here, we identify adult stem cells and their differentiating progeny as additional target tissues for ApoB-mediated NL transport in a regeneration competent animal, and propose utilizing planarians as models to understand the influence of lipid metabolism on stem cells during regeneration.

## Results

### Planarian *apolipoprotein b* orthologs are expressed by intestinal cells

Previously, we demonstrated that knockdown of an intestine-enriched transcription factor, *nkx2.2*, inhibited neoblast proliferation and formation of the regeneration blastema, the unpigmented mass of tissue produced after amputation^39^. These observations suggested that the intestine could play a non-autonomous role in regulating stem cell dynamics. In an effort to identify intestine-enriched transcripts encoding neoblast regulators, we generated transcriptomes from control and *nkx2.2(RNAi)* planarians, in which neoblast proliferation and regenerative tissue production are severely inhibited (Supplementary Fig. 1a-c)^39^. An ortholog of human *apolipoprotein b* (“*apob-1*”) encoding conserved Vitellogenin, DUF1943, and von Willebrand Factor D domains, was the second-most-downregulated transcript by significance and fold-change, while a paralog, *apob-2,* was also significantly downregulated (Fig. 1a; Supplementary Fig. 1d, e; Supplementary Data 1a-c; and Supplementary Data 2a-b). Consistent with single cell expression profiling^40^ (Supplementary Fig. 1f), both transcripts were highly enriched in intestinal phagocytes, absorptive cells responsible for digestion, nutrient storage, and metabolite secretion (Fig. 1b and Supplementary Fig. 1f, g). In addition, both transcripts were weakly expressed in a small number of cells outside the intestine likely to be differentiating neoblast progeny (Fig. 1b and Supplementary Fig. 1f).

**Figure 1.**
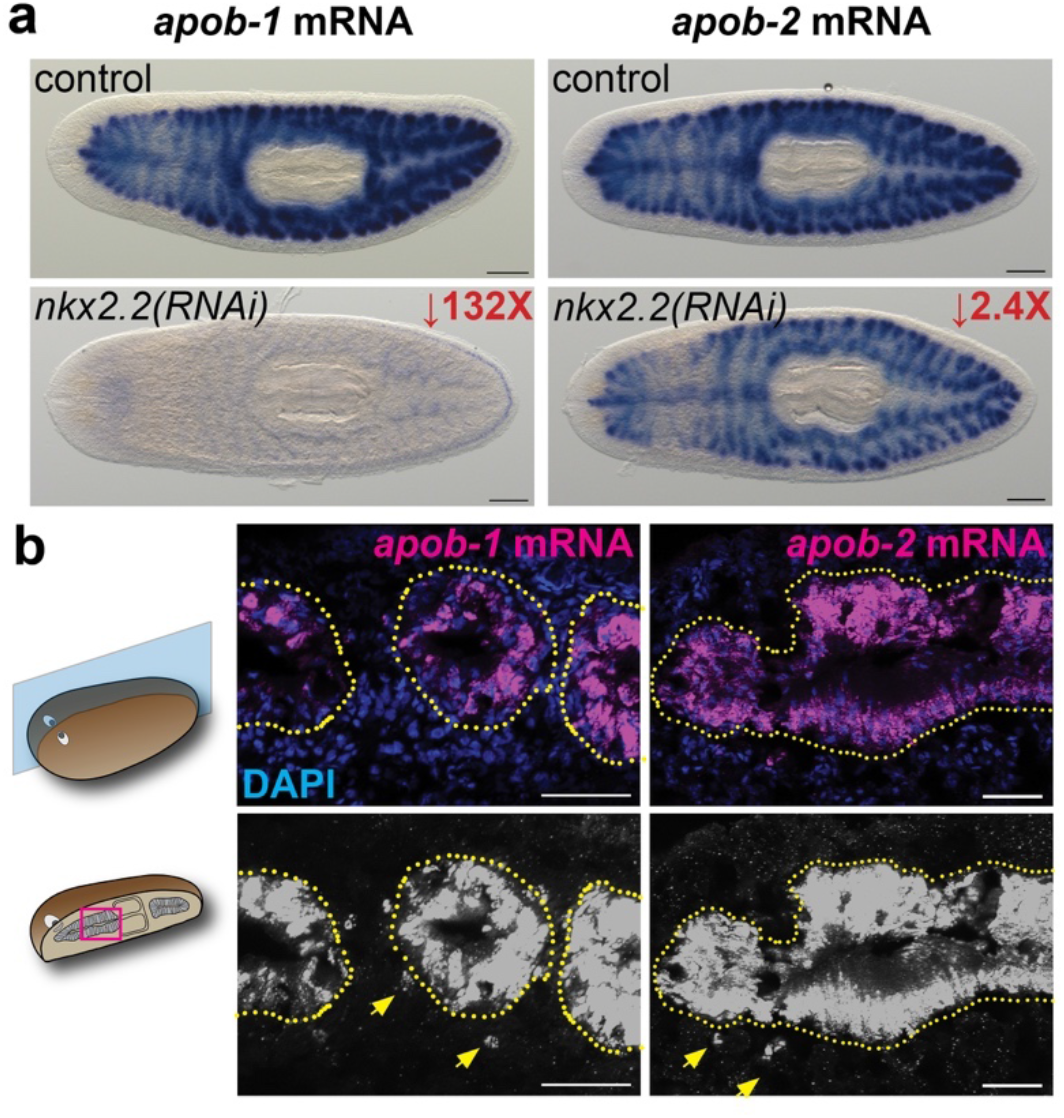
Transcripts encoding planarian *apolipoprotein b* orthologs are intestine-enriched and downregulated in *nkx2.2(RNAi)* animals. **(a)** *apob-1* and *apob-2* mRNA (ISH) expression in control (top) and *nkx2.2(RNAi)* planarians. **(b)** *apob-1* and *apob-2* mRNA expression (FISH) in sagittal sections. Arrows indicate *apob*-expressing cells (likely differentiating phagocytes) outside the intestine (dotted yellow outline) in digitally brightened images. Scale bars: 200 µm **(a)**; 50 µm **(b)**.

### ApoB orthologs are required for viability and regulate neutral lipid transport

To determine whether ApoB orthologs were required for homeostatic tissue renewal and/or regeneration, we knocked down *apob-1* and *apob-2* using RNA interference^41^. In uninjured planarians, knockdown of either *apob-1* or *apob-2* individually had no phenotype (not shown), suggesting functional redundancy. However, after 3-5 double-stranded RNA (dsRNA) feedings, double knockdown *apob(RNAi)* animals displayed phenotypes that progressed from mild (modest/regional pigmentation loss), to severe (animal-wide pigmentation loss and reduced motility), to very severe (head regression, ventral curling, and eventual lysis) (Fig. 2a). Head regression and ventral curling phenotypes are common in planarians lacking functional neoblasts^42^, and suggested that ApoB orthologs could have functional roles in the regulation of planarian stem cells.

**Figure 2.**
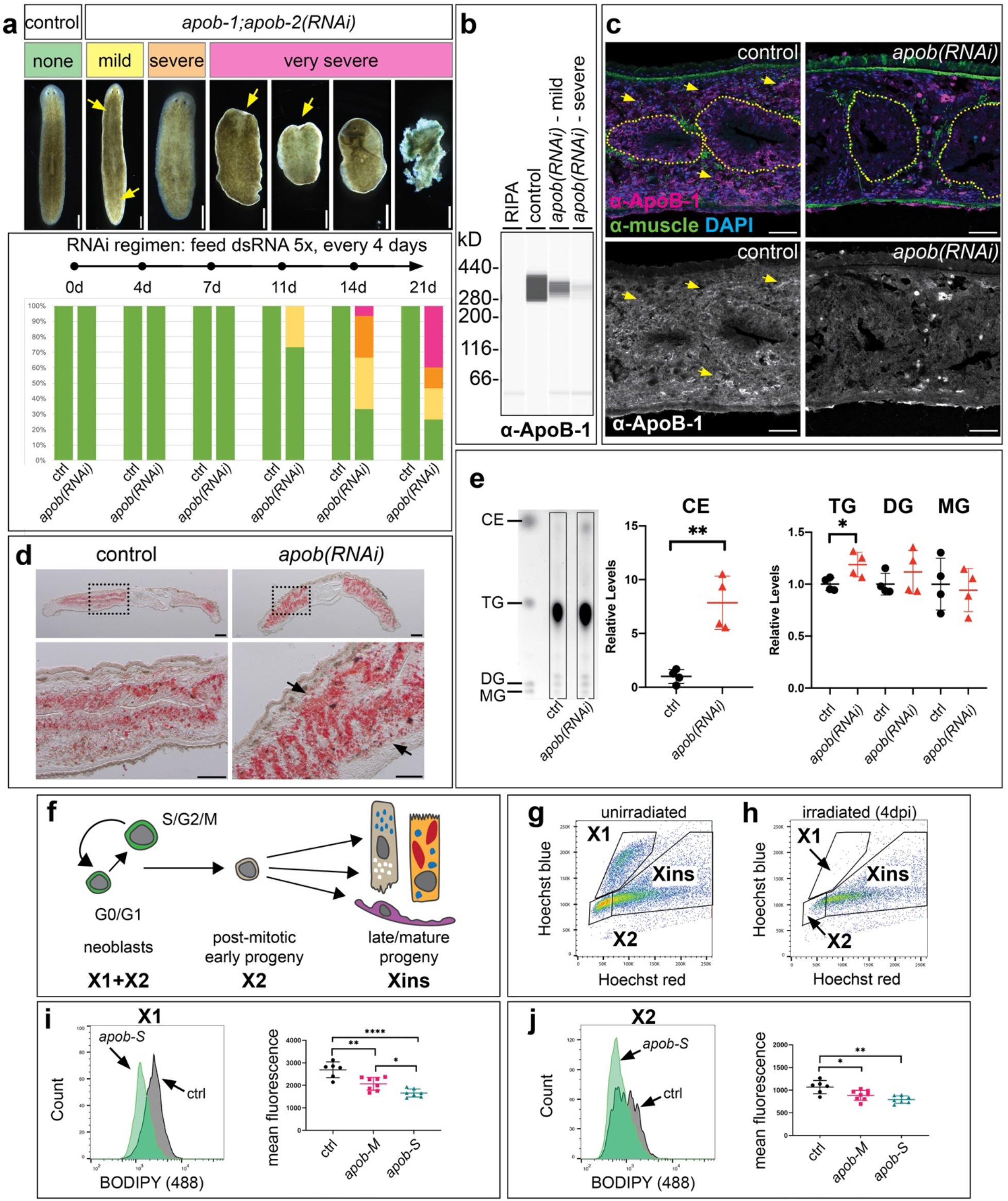
*apob* orthologs are required for viability and neutral lipid metabolism. **(a)** Simultaneous RNAi-mediated knockdown of *apob-1* and *apob-2* caused mild (yellow) and severe (orange) depigmentation, and very severe (pink) phenotypes including head regression, ventral curling, and lysis. Knockdown was initiated with n = 15 animals per condition. **(b)** Simple Western (capillary-based protein analysis) lane view of extracts from control and knockdown planarians labeled with custom anti-ApoB-1. **(c)** ApoB-1 protein expression in sagittal sections. Expression in intestine (dotted outline) and outside the intestine (arrows) was dramatically reduced in “very severe” *apob-1(RNAi);apob-2(RNAi)* planarians (right panels). mAb 6G10 (green) labeled body wall, dorsoventral, and visceral muscle fibers (demarcating the intestine.) **(d)** Neutral lipids accumulated in the intestine and parenchyma (arrows) of “mild” *apob(RNAi)* planarians. Oil Red O labeling, sagittal sections. **(e)** Cholesteryl esters (CE) and triglycerides (TG), but not diacylglycerides (DG) or monoglycerides (MG) were significantly elevated in “mild” *apob(RNAi)* animals. Thin layer chromatography, intensities of lipid species measured in ImageJ and normalized to controls. Student’s t test (unpaired, two-tailed); *p*=.0017 (CE) and *p*=.0303 (TG). Error bars: mean ± S.D., n=4. **(f)** Lineage schematic indicating cell types in X1, X2, and Xins subpopulations. **(g)** Example flow plot for uninjured animals. **(h)** Example plot for animals 4 days post irradiation (“4 dpi”), showing ablation of cells in X1 and depletion of cells in X2. **(i-j)** *apob* knockdown caused reduction of neutral lipids in X1 and X2 cells. One-way ANOVA with Tukey’s multiple comparisons test. Error bars: mean ± S.D., n≥6 biological replicates per condition. Scale bars: 50 µm **(c)**; 200 µm **(d,** upper panels); 100 µm **(d,** lower panels).

ApoB orthologs in vertebrates and insects regulate LP biogenesis and secretion^5, 43^. To test whether ApoB protein was secreted, we made a custom antibody recognizing the N-terminus of Smed-ApoB-1 (Fig. 2b, c). ApoB-1 protein was enriched in the intestine, but also throughout the parenchyma (tissues surrounding other organs, sandwiched between the epidermis and intestine, where neoblasts also reside). This indicated that although *apob* mRNAs were intestine-enriched (Fig. 1a, b), ApoB protein was robustly secreted and transported to peripheral tissues (Fig. 2c). Expression was dramatically reduced in both regions in *apob-1;apob-2(RNAi)* double knockdown planarians (*“apob(RNAi)”* hereafter for brevity), demonstrating that knockdown effectively reduced ApoB protein levels (Fig. 2c).

ApoB orthologs facilitate NL secretion and transport via LPs^5, 38^, while ApoB binding by receptors mediate NL uptake and metabolism by target cells^44^. To test whether planarian ApoB functioned similarly, we first evaluated NL distribution in *apob(RNAi)* animals. In histological sections, NLs (labeled with Oil Red O) were elevated in both the intestine as well as tissues surrounding the intestine, suggesting that both LP secretion by the intestine and uptake/metabolism by peripheral tissues were compromised (Fig. 2d). Using thin layer chromatography, we also found significant elevation of cholesteryl esters and triglycerides in lipid extracts from *apob(RNAi)* animals (Fig. 2e). These phenotypes predicted that delivery of NLs to neoblasts and/or their progeny would be compromised by ApoB inhibition. To test this, we quantified NL content in neoblasts and their progeny using a fluorescent NL probe, BODIPY-493/503, by flow cytometry. Three major subpopulations of planarian cells can be distinguished by their DNA content and sensitivity to X-irradiation^45, 46^ (Fig. 2f-h). The “X1” fraction/gate includes >2C DNA content neoblasts in S/G2/M phase of the cell cycle, while “X2” includes 2C neoblasts in G0/G1 and G0 post-mitotic progeny. “Xins,” named for the fact that cells in this gate are insensitive to X-irradiation, consists of later stage progeny and mature differentiated cells. We found a dramatic reduction of fluorescence in both the X1 and X2 fractions in *apob(RNAi)* animals vs. controls (Fig. 2i, j), indicating that a reduction of NL content in neoblasts and their progeny was caused by ApoB inhibition. Together, these data demonstrated that planarian ApoB proteins were produced by the intestine and were likely secreted as LPs to transport NLs from the intestine to stem cells and their differentiating progeny.

### *apob* and *lipoprotein receptor* genes are upregulated in regenerating fragments

Amputation induces changes in gene expression during planarian regeneration^47–49^. We predicted that if *apob* and other genes involved in neutral lipid metabolism played functional roles during regeneration, they would be up- or down-regulated in amputated tissue fragments. Using quantitative PCR, we found that both *apob-1* and *apob-2* transcripts were upregulated in tissue fragments at two distinct time points during earlier stages of regeneration commonly associated with neoblast proliferation (1-3 days) and differentiation (4-5 days) (Supplementary Fig. 2a, b). Upregulation of *apob-1* and *apob-2* was also observed in previously published RNA-Seq data from a 14-day time course of whole-body planarian regeneration (Supplementary Fig. 2c)^49^. Using quantitative capillary-based Western blotting, we also found that ApoB-1 protein levels increased by 3-5 days after amputation (Supplementary Fig. 2d). Consistently, accumulation of neutral lipids was sustained in 3- and 6-day *apob(RNAi)* regenerates (Supplementary Fig. 2e, f), suggesting that LP trafficking from the intestine was disrupted, as in uninjured animals (Fig. 2d).

In addition, we found that three planarian orthologs of human lipoprotein receptors (which bind apolipoproteins, enabling LP uptake/metabolism) were upregulated in the blastema, and regenerating brain and pharynx (Supplementary Fig. 3a, b, d). These transcripts were expressed in both *piwi-1-*mRNA-positive neoblasts as well as *piwi-1-*negative cells that were likely to be differentiating neoblast progeny in the regenerating pharynx and blastema (Supplementary Fig. 3d). Two lipoprotein receptor genes (*ldlr-1* and *ldlr-2*) were also differentially expressed in the whole-body regeneration RNA-Seq dataset (Supplementary Fig. 3c)^49^. Using the Gene Ontology to mine the same dataset^49^, we identified additional planarian genes predicted to regulate lipid transport, triglyceride, and cholesterol metabolism, and found that many of these transcripts were also up- and down-regulated during regeneration (Supplementary Fig. 3e and Supplementary Data 3). Together, these observations indicate that *apob-1, apob-2,* and other genes involved in neutral lipid metabolism were differentially expressed in response to amputation, consistent with roles during regeneration.

### *apob* paralogs are required for polarity re-establishment and organogenesis during regeneration

To assess functional roles of *apob* paralogs during regeneration (Fig. 3a), we chose uninjured animals after three dsRNA feedings that exhibited “mild” and “severe” phenotypes (Fig. 2a), amputated their heads, and assessed blastema morphogenesis, which absolutely requires neoblast proliferation and differentiation^50–52^. Controls, *apob-1(RNAi)-*only, and *apob-2(RNAi)-*only fragments regenerated normally (Fig. 3b), again suggesting *apob* paralogs likely functioned redundantly. However, *apob(RNAi)* double knockdown animals had reduced posterior blastemas (head fragments) and anterior blastemas (trunk fragments) at 8 days post-amputation (Fig. 3b). Blastema size was noticeably smaller in fragments from “severe” (“*apob-S”*) animals compared to “mild” (“*apob-M”*) animals (Fig. 3b), suggesting that neoblasts and/or their progeny were more strongly affected over time.

**Figure 3.**
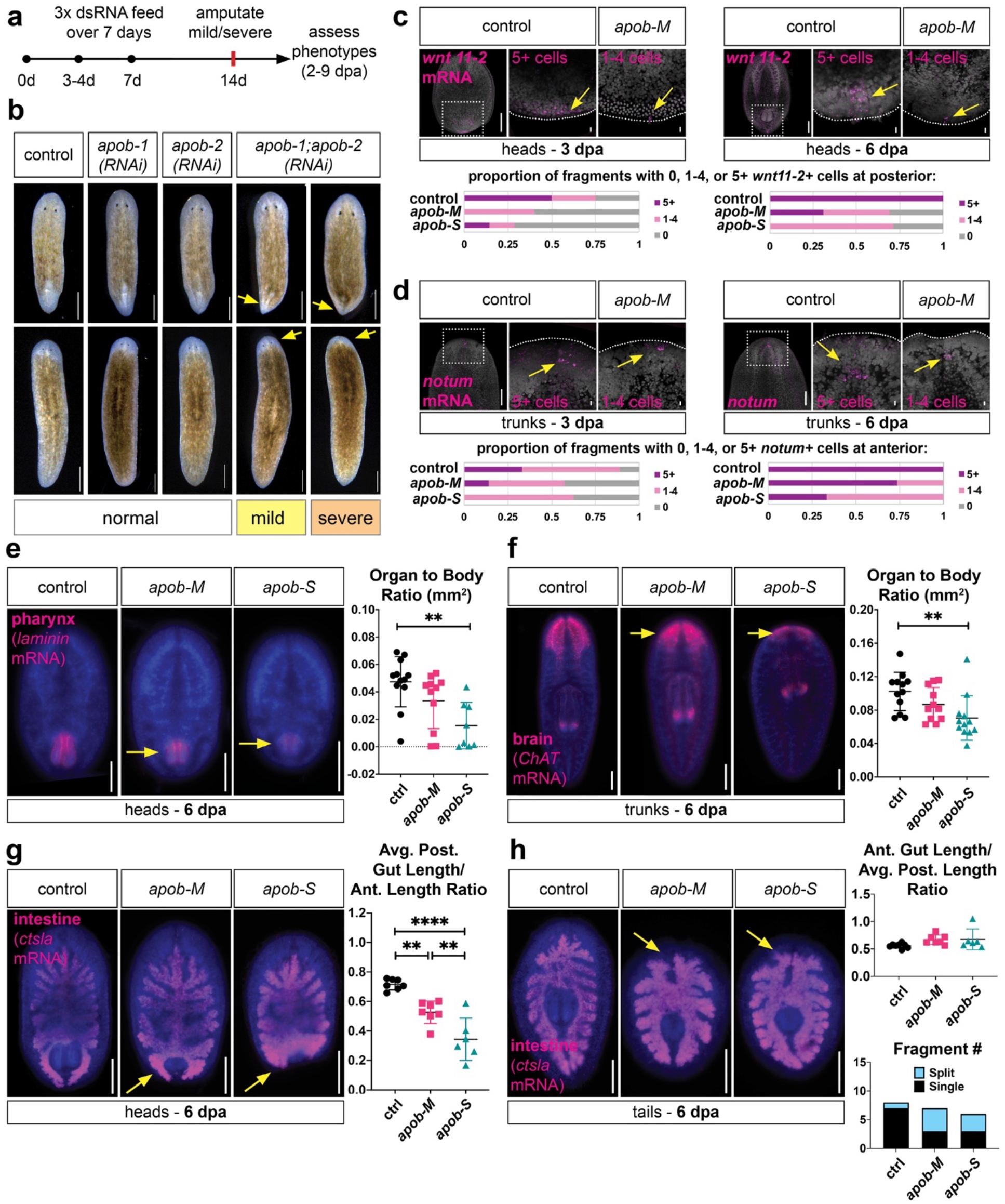
*apob* inhibition delays regeneration. **(a)** Schematic of RNAi treatment and analysis regimen. **(b)** *apob* double knockdown resulted in smaller blastemas (arrows) in head and trunk regenerates (8 dpa). **(c-d)** Differentiation of cells expressing polarity genes *wnt11-2* (posterior) and *notum* (anterior) was delayed in *apob(RNAi) animals.* FISH; max projections of confocal images. n=5-15 fragments per condition were scored for number of positive cells. Control and *apob-M* fragments shown as phenotype examples. **(e)** Pharynx regeneration was reduced in *apob(RNAi)* head regenerates (6 dpa) (*laminin* mRNA FISH, epifluorescent images). p=.0021, control vs. *apob-S.* **(f)** CNS regeneration was reduced in *apob(RNAi)* tail regenerates (6 dpa) (*ChAT* mRNA FISH, epifluorescent images). p=.0101, control vs. *apob-S.* **(g-h)** *apob* RNAi disrupted intestine regeneration (6 dpa) (*ctsla* mRNA FISH, epifluorescent images). New branches were shorter in heads **(g)**. p=.0004 (*apob-M*); p=.0016 (*apob-S*). New branches often failed to fuse in tails **(h)**. Significance testing in **(e-h)**: one-way ANOVA with Tukey’s multiple comparisons test. Error bars: mean +/- S.D., n≥7 per condition. DAPI (blue) in **(e-h)**. Scale bars: 500 µm **(b)**; 200 µm **(c-d);** 10 µm **(c-d** insets); 200 µm **(e-h)**.

Expression of so-called “position control genes” to re-establish axial polarity is an essential early event required for whole-body regeneration^53^. We asked whether expression of anterior (*notum*)^54^ or posterior (*wnt11-2*)^55^ was affected in *apob(RNAi)* regenerates. Indeed, at three days post-amputation (3 dpa), the number of both posterior *wnt11-2-*expressing cells (Fig. 3c) and anterior *notum*-expressing cells (Fig. 3d) was reduced in *apob-M* and *apob-S* fragments. By 6 dpa, although most *apob(RNAi)* fragments had regenerated more of these cells, many fragments (especially *apob-S*) still had fewer than five cells compared to control fragments (Fig. 3c, d). These data suggested that *apob* inhibition delayed, but did not completely abolish, re-establishment of these cells.

*apob* RNAi also affected regeneration of planarian organs, including the brain, pharynx, and intestine. Although both brain and pharynx regenerated, these organs were smaller than in controls (Supplementary Fig. 2g, h). This suggesting regeneration was delayed, but not blocked, similar to the phenotype for cells expressing polarity cues. Quantitatively, at 6 dpa, both brain and pharynx were smaller in size, especially in *apob-S* fragments (Fig. 3e, f). To assess intestine, we analyzed both head and tail regenerates, in which a combination of neoblast-driven new cell production and remodeling of pre-existing, differentiated intestinal branches is required for successful regeneration^56^. Newly regenerated posterior branches were significantly shorter in *apob(RNAi)* head fragments (Fig. 3g). By contrast, in tail fragments, length of the new anterior intestine was not significantly affected (Fig. 3h). Rather, these branches failed to fuse at the anterior midline, leading to a “split” anterior branch phenotype in ∼50% of *apob(RNAi)* tail fragments (Fig. 3h). Together, these results suggested that ApoB reduction delayed regeneration of multiple cell types and organs, and, in the case of the intestine, affected differentiation of new cells as well as collective cell migration or other processes required for remodeling.

### *apob* knockdown causes accumulation of early neoblast progeny

Because *apob* knockdown disrupted multiple neoblast-driven processes during regeneration, we asked whether defects in neoblast maintenance or proliferation were responsible for the effects of *apob* RNAi on regeneration. First, we assessed neoblast numbers by FISH using *piwi-1,* a pan-neoblast marker, and *tgs-1,* a marker for a more pluripotent subpopulation (which also includes neural specialized neoblasts)^49, 50, 57^. In both uninjured planarians and 7.5-day regenerates, neoblast numbers and distribution were grossly normal (Supplementary Fig. 4a, b). We also assessed proliferation using anti-phospho-Histone H3 (Ser10) (“PS10”) immunolabeling, which marks cells in late G2 and M phase of the cell cycle^58, 59^. In uninjured animals and 2-day head regenerates, the number of mitotic neoblasts increased modestly in *apob-M,* but not *apob-S* samples (Fig. 4a, b). In 2-day trunk regenerates, there was a more significant (∼50%) increase in *apob-M* fragments, and a modest increase in *apob-S* fragments (Fig. 4b). Together, these results suggested that ApoB reduction might cause moderate hyperproliferation, or, alternatively, a modest mitotic delay, without dramatically affecting *piwi-1+* or *tgs-1+* neoblast numbers. These mild phenotypes also raised the possibility that ApoB reduction might preferentially dysregulate differentiation, rather than proliferation or maintenance of actively cycling neoblasts.

**Figure 4.**
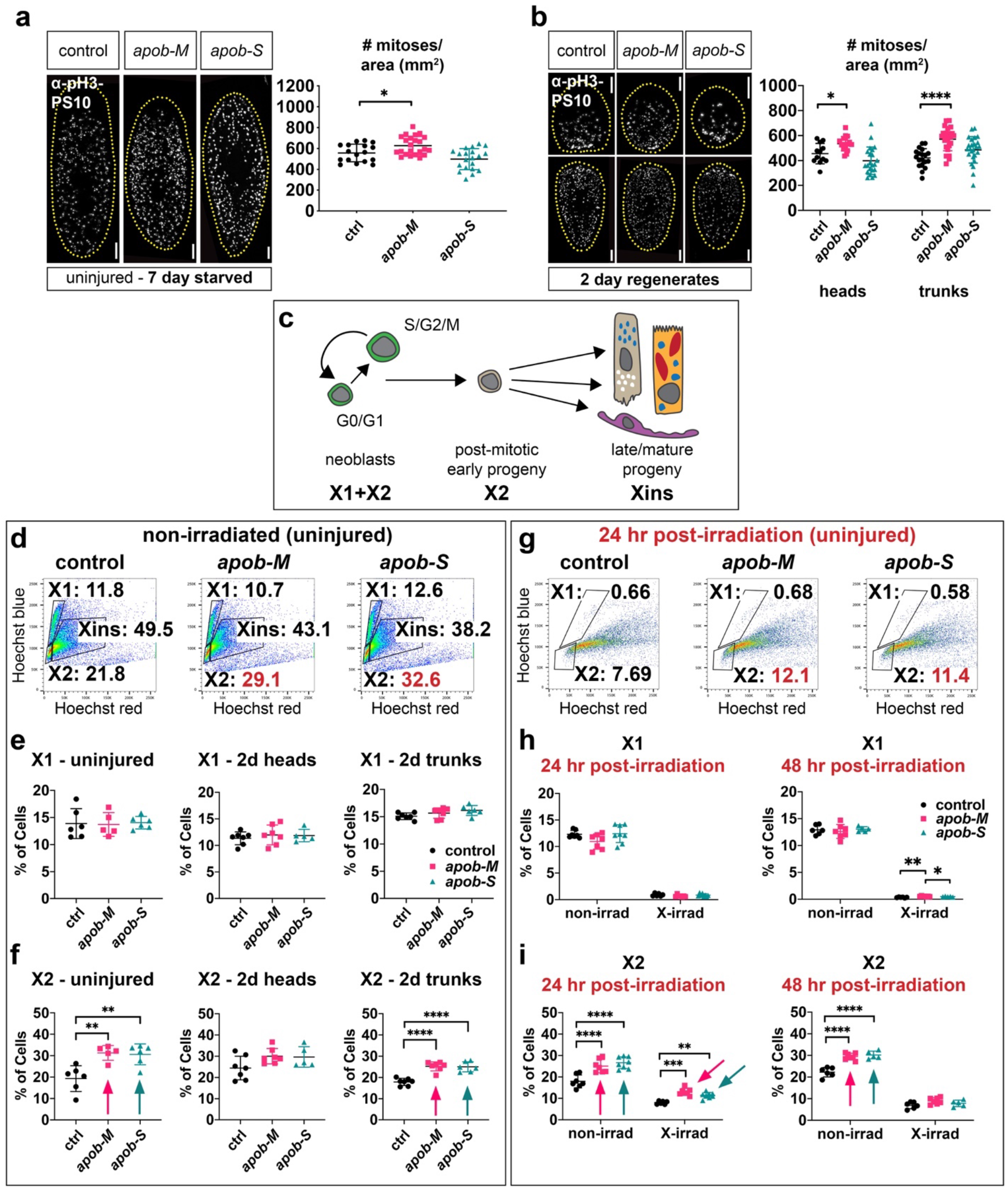
*apob* inhibition causes accumulation of neoblast progeny. **(a-b)** PhosphoHistone-H3-S10-positive (pH3-PS10) cells were modestly elevated in *apob-M,* but not *apob-S,* uninjured animals **(a)** and 2-day regenerates **(b)**. One-way ANOVA with Dunnett’s T3 multiple comparisons test. Error bars: mean +/- S.D., n≥12 animals/fragments per condition **(a-b)**. **(c)** Lineage schematic indicating cell types/states in X1, X2, and Xins subpopulations. **(d)** Examples of flow dot plots from uninjured planarians indicating percentages of cells in each gate, with X2 increase in *apob(RNAi)* animals in red. **(e-f)** Percentage of cells in X1 **(e)** and X2 **(f)** in uninjured (left), 2-day head (middle), and 2-day trunk (right) regenerates. Arrows indicate statistically significant increases in X2. **(g)** Examples of flow plots from uninjured planarians, 24 hr post-irradiation, indicating percentages of cells in each gate, with X2 increase in *apob(RNAi)* animals in red. **(h-i)** Percentage of cells in X1 **(h)** and X2 **(i)** in uninjured planarians, 24 hr (left) and 48 hr (right) post-irradiation. Arrows indicate significant increases in X2. One-way ANOVA and Tukey’s multiple comparisons test for non-irradiated samples and X1 irradiated samples (*p* values in **e-f, h**, see Methods). Two-way ANOVA and two-stage linear step-up procedure of Benjamini, Krieger and Yekutieli for multiple comparisons for X2 irradiated samples (*q* values in **i**). Although differences between RNAi conditions in X1 at 48 hr post-irradiation were significant, percent of cells in this gate was negligible (<0.7% in all samples). Error bars = mean ± S.D, n≥5 biological replicates per condition. Scale bars: 200 µm **(a-b)**.

We tested these possibilities quantitatively using flow cytometry. We dissociated uninjured and regenerating planarians, and evaluated the proportions of cells in the X1 and X2 fractions (Fig. 4c). In uninjured planarians, as well as 2-day and 7-day regeneration fragments, X1 neoblast numbers were unaffected by *apob* knockdown (Fig. 4d, e and Supplementary Fig. 4c), consistent with the moderate effect of *apob* RNAi on neoblast proliferation (Fig. 4a, b and Supplementary Fig. 4a, b). By contrast, in uninjured planarians as well as 2- and 7-day trunk fragments, the proportion of cells in the X2 gate increased significantly by ∼20-40% (Fig. 4f and Supplementary Fig. 4c) in *apob(RNAi)* animals vs. control. In 2 dpa head fragments, there was also a modest, but statistically insignificant, increase (∼15%) in the X2 fraction (Fig. 4f). Because the X2 fraction includes both cycling G1-phase neoblasts and differentiating post-mitotic progeny^60, 61^, these results suggested that *apob* inhibition might cause lengthening of G1 phase of the cell cycle, and/or a delay in differentiation of neoblast progeny, either of which could increase the proportion of cells in X2.

In order to distinguish between these possibilities, we examined the X2 fraction in uninjured planarians 24 hr after X-irradiation, which preferentially ablates over 95% radiation-sensitive cycling neoblasts, without affecting many early neoblast progeny^49, 52, 60^. As expected, the X1 fraction was almost completely eliminated in both control and *apob(RNAi)* samples (Fig. 4g, h). However, although irradiation reduced the proportion of X2 cells by ∼48-56% in both control and *apob(RNAi)* animals (Fig. 4i), the expansion of the X2 fraction persisted in *apob-M* (∼62.5% increase) and *apob-S* animals (∼42.5% increase) relative to controls (Fig. 4i). This increase in radiation-insensitive cells in X2 strongly suggested that the primary defect in *apob* knockdown animals was a delay in differentiation of neoblast progeny.

Intriguingly, the increase in radiation-resistant X2 cells was transient: by 48 hr post-irradiation, the proportion of X2 cells in *apob(RNAi)* samples was only modestly (but not significantly) elevated (Fig. 4i). This suggested that differentiation was delayed, but not arrested, in *apob(RNAi)* animals. One possible explanation for this observation is that, at later time points, progeny had differentiated further, and resided in the Xins gate (rather than the X2 gate) as they achieved a more mature cell state. Alternatively, some early progeny might have undergone apoptosis or necrosis at later time points, reducing the apparent accumulation of neoblast progeny in X2.

### ApoB reduction preferentially dysregulates expression of transcripts enriched in differentiating neoblast progeny and mature cell types

Our flow cytometry data suggested that the primary phenotype in uninjured and regenerating *apob(RNAi*) animals was a delay in the differentiation of neoblast progeny. To test this interpretation further, we performed whole animal RNA sequencing on control, *apob-M,* and *apob-S* animals, and identified differentially expressed (DE) genes in the two *apob(RNAi)* groups (Supplementary Fig. 5a-c and Supplementary Data 4). Then, to identify specific biological processes affected by *apob* inhibition, we used the Gene Ontology (GO) to identify patterns of dysregulation in specific functional categories (Supplementary Fig. 5d, e and Supplementary Data 5). In addition, to determine whether *apob* knockdown disproportionately dysregulated genes enriched in differentiating progeny states, we mapped the global *apob* “dysregulation signature” to published bulk- and single-cell transcriptomes to determine which cell types and states were most affected by *apob* inhibition.

*apob* knockdown dysregulated thousands of genes, causing upregulation of 842 (*apob-* M) and 1960 (*apob-S)* transcripts, and downregulation of 1139 (*apob-*M) and 2547 *(apob-*S) transcripts, relative to control (Supplementary Fig. 5a-c and Supplementary Data 4a-d). For upregulated transcripts, “lipid metabolism” was the fifth-most over-represented Biological Process (BP) term in *apob-M* animals, and was the most over-represented category in *apob-S* animals (Supplementary Fig. 5d and Supplementary Data 5a, b). Enrichment of the subcategories of acylglycerol, fatty acid, steroid, and glycerolipid metabolism was consistent with known roles of ApoB orthologs, and suggested a possible compensatory gene expression response to *apob(RNAi)-*induced disruption of NL transport. *apob(RNAi)* also downregulated transcripts in additional metabolism categories, including gluconeogenesis, glycolysis, pyruvate, and nucleotide metabolism (e.g., NADH and ADP), as well as ion transport, indicating wide-ranging dysregulation of metabolite processing and trafficking, especially in *apob-S* animals (Supplementary Fig. 5e and Supplementary Data 5c, d). Interestingly, *apob* inhibition dysregulated many additional, non-metabolism-related transcripts. Upregulated functional categories included differentiation and/or specification of multiple tissues including epidermis, nervous system, and the eye, while downregulated categories included cilium morphogenesis and function (possibly reflecting disruption of ciliated epidermal and/or protonephridial cell numbers or physiology), muscle morphogenesis and function, and extracellular matrix organization (Supplementary Fig. 5d, e and Supplementary Data 5a-d). Importantly, categories related to cell cycle, mitosis, etc. were not enriched among up- or downregulated transcripts. Together, these data suggested that ApoB reduction affected metabolism, cell/tissue differentiation, and functions of mature cell types, but not processes required specifically for cellular proliferation.

Next, to test whether *apob* disproportionately dysregulated genes expressed by differentiating progeny and mature cells, we cross-referenced our DE transcript list with recently published bulk transcriptomes from flow-sorted X1, X2, and Xins cell fractions (Fig. 4c)^49^, as well as cells sorted based on their expression of PIWI-1 protein, a widely used marker whose expression marks states within planarian cell lineages (e.g., high PIWI-1 levels in neoblasts, low PIWI-1 levels in some neoblasts and differentiating progeny, and negligible PIWI-1 levels in mature cells)^49^. We first identified transcripts that were enriched in each cell fraction to generate a “signature” transcript list for each fraction (Supplementary Fig. 6a, c). We then asked what percent of these signature transcripts were dysregulated in *apob-M* and *apob-S* animals (Fig. 5a, b and Supplementary Fig. 6b, d). *apob* RNAi dysregulated signature transcripts in all six fractions. However, compared to X1 or “PIWI-HI” signature transcripts, 2-4 times as many transcripts were up- or down-regulated in X2 and “PIWI-LO” cell classes, which included both neoblasts and early progeny (Fig. 5a, b). Relative to Xins and PIWI-NEG fractions, there were also more dysregulated transcripts represented in the X2 and PIWI-LO fractions in *apob* RNAi animals (Fig. 5a, b). To demonstrate the validity of this approach, we cross-referenced transcriptomes from planarians 24 hours after irradiation^49^, and found that dysregulated transcripts overlapped mainly with X1 and PIWI-HI signature transcripts, and were primarily downregulated, consistent with neoblast loss (Supplementary Fig. 6e, f). Together, this analysis demonstrated that *apob* knockdown preferentially affected transcripts enriched in differentiating progeny, an observation that is consistent with accumulation of cells in this state in flow cytometry experiments (Fig. 4d-i).

**Figure 5.**
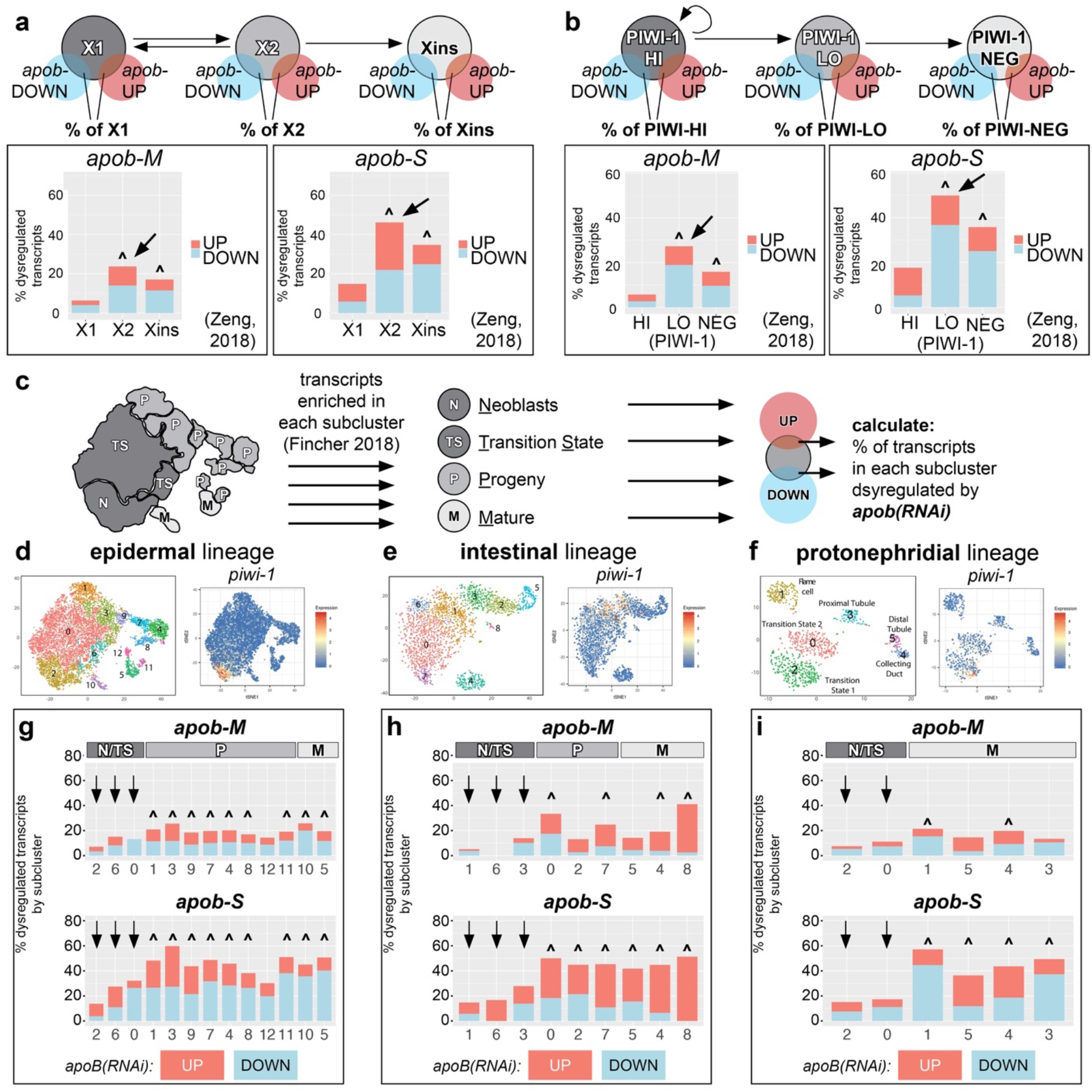
*apob* RNAi preferentially dysregulates transcripts in differentiating neoblast progeny and mature cell states. **(a-b)** *apob* RNAi dysregulates greater proportions of X2 **(a)** and PIWI-LO **(b)** signature transcripts (arrows in histograms). Venn diagrams at top show analysis scheme: percent of X1/X2/Xins **(a)** and PIWI-HI/PIWI-LO/PIWI-NEG **(b)** signature transcripts ^49^ that overlap with transcripts dysregulated in *apob-M/apob-S* animals (this study) (see also Supp. Fig. 6a-d). Histograms show percentage of signature transcripts up- and down-regulated in *apob-M/apob-S* animals. **(c)** Schematic example (for epidermal lineage) illustrating how transcripts dysregulated in *apob(RNAi)* animals were cross-referenced with neoblast (N), transition state (TS), progeny (P), and mature (M) cell state subclusters from^40^. See Methods for details. **(d-f)** t-SNE plots (digiworm.wi.mit.edu) indicate subclusters and *piwi-1* mRNA expression for each lineage. **(g-i)** *apob* knockdown dysregulated greater proportions of transcripts enriched in progeny (“P”) and mature (“M”) cell subclusters in multiple cell type lineages. Arrows indicate less-affected transcripts enriched in neoblasts/transition state (“N/TS”) subclusters. Carets (^) **(a, b, g-i)** indicate significant gene expression overlap (*p<*0.05, Fisher’s exact test, see Source Data 2 for individual *p* values).

To determine whether *apob* RNAi preferentially affected specific lineages or states within individual lineages, we also compared the gene expression signature of *apob(RNAi)* animals with recently published single cell transcriptomes^40^. Specifically, we cross-referenced transcripts dysregulated by *apob* RNAi with transcripts enriched in cell state subclusters in eight planarian lineages (Fig. 5c-i, Supplementary Fig. 8a, and Supplementary Fig. 8c-l). The resulting “dysregulation signature” for each lineage showed how *apob* RNAi affected gene expression in specific cell states during the progression from pluripotent cycling neoblasts (“N”), to early transition states (“TS”), to differentiating progeny (“P”) to mature cell states (“M”) (see example schematic for epidermal lineage in Fig. 5c). We observed a striking and consistent dysregulation pattern in all eight lineages. In three well characterized lineages (epidermis, intestine, and protonephridia) (Fig. 5c-i), the percentage of dysregulated “state-enriched” transcripts was lowest for neoblast and early transition state subclusters, but higher for progeny and mature cell states. Similar trends were observed for less characterized lineages such as muscle, pharynx, and *cathepsin*-positive cells (Supplementary Fig. 8c-l). Validating this approach, irradiation^49^ primarily dysregulated neoblast and transition state transcripts (Supplementary Fig. 7a-g), and, to a lesser degree for some lineages, progeny-enriched transcripts (Supplementary Fig. 8a and Supplementary Fig. 8m-q). Similarly, and as expected, X1- and PIWI-HI-enriched transcripts overlapped with neoblast and transition state transcripts, while X2-/PIWI-LO and Xins/PIWI-NEG transcripts were enriched in progressively later cell state subclusters in each lineage (Supplementary Fig. 7h-k, Supplementary Fig. 8b, and Supplementary Fig. 8r-v). Together, these results provided further evidence that *apob* inhibition specifically dysregulated gene expression in progeny and mature cell types/states in all eight lineages, and lent additional support to the conclusion that ApoB was required for differentiation of planarian neoblasts.

## Discussion

In this study, we discover a non-cell autonomous role for ApoB proteins in regulation of planarian neoblast differentiation. Our data suggest that ApoB transports NLs from the intestine to both neoblasts and their differentiating progeny via LPs (Fig. 6). In the absence of ApoB, neoblast proliferation and maintenance are largely unaffected, but differentiation is delayed, causing accumulation of differentiating progeny (Fig. 6) and slowing regeneration.

**Figure 6.**
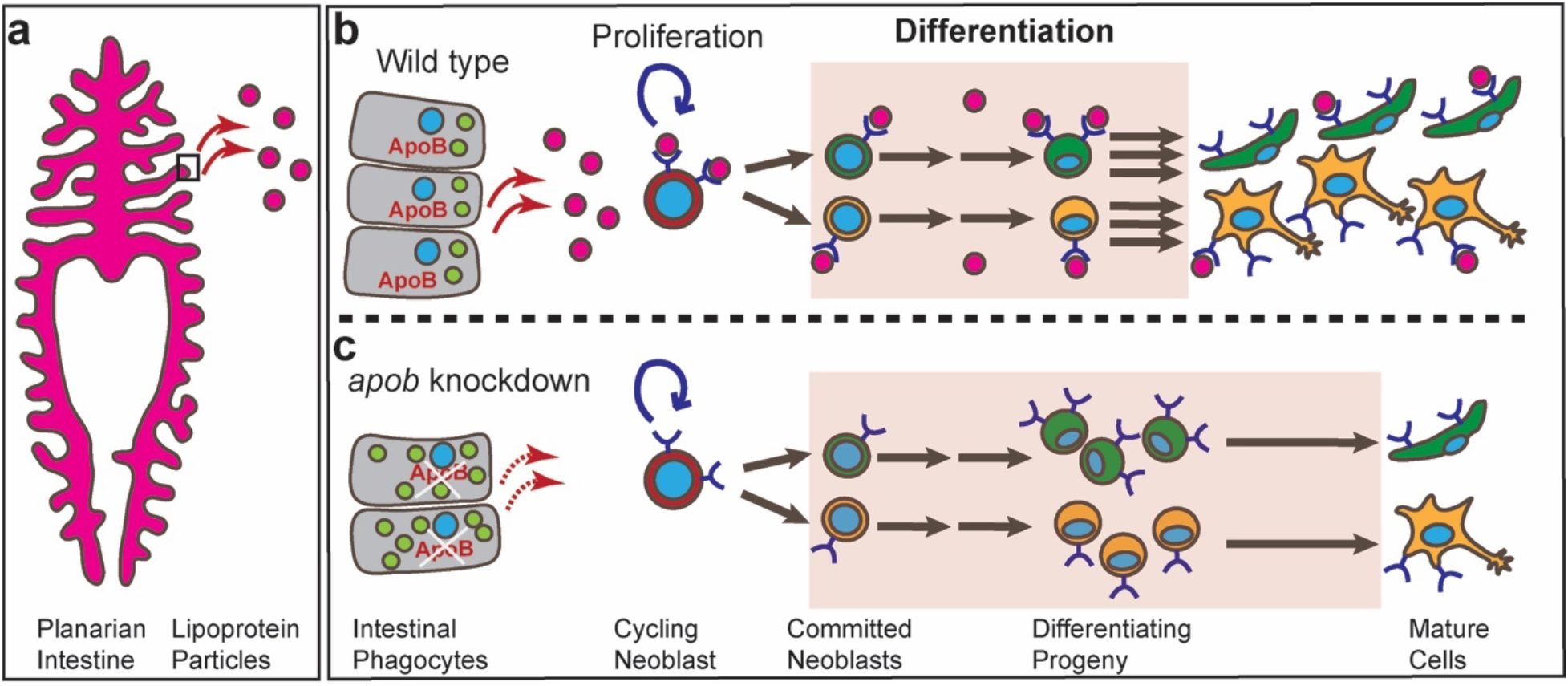
A putative model for ApoB function in regulating differentiation of planarian stem cell progeny. **(a)** Our data support a working model in which ApoB is expressed by phagocytes in the intestine, a primary site of LP production and secretion. **(b)** ApoB mediates secretion of neutral lipids in LPs from intestinal phagocytes to neoblasts and their progeny. **(c)** In the absence of ApoB, lipids accumulate in the intestine, and LP delivery to neoblasts and their progeny is disrupted, reducing their neutral lipid content. Neoblast proliferation and renewal are largely unaffected by reduced ApoB function. Instead, differentiation and later maturation of most, if not all, planarian cell lineages are slowed, causing an accumulation of differentiating progeny and a delay in regeneration of multiple organs. Box in **(a)** represents region magnified in **(b)** wild type and **(c)** *apob* knockdown conditions. Nuclei, blue. Lipid droplets, light green. LPs, magenta. Apical/lumenal phagocyte surface is to the left and basal/mesenchymal phagocyte surface is to the right in **(b, c)**.

*apob* knockdown causes a ∼40% increase in a flow cytometry fraction of 2C DNA content cells (X2) that includes both cycling neoblasts in G1, and post-mitotic differentiating progeny in G0^57, 60^. This increase occurs without simultaneously affecting the >2C S/G2/M cell fraction (X1), and is sustained even in recently irradiated animals, indicating that the expansion is comprised primarily of G0 progeny, and not G1 neoblasts. We suggest that this phenotype indicates a disruption in the progression of differentiation after cell fate specification, and not specification itself. In planarians, specification likely begins during S-phase, because expression of fate-specific transcription factors is significantly higher in S/G2/M neoblasts than in G1 neoblasts^57, 62^. Intriguingly, inhibition of other planarian genes required for differentiation (e.g., the transcription factor *mex3-1,* the extracellular matrix component *collagen4-1,* and the transcriptional co-activating protein *cbp-3*) also cause increases in neoblast numbers *in vivo*^52, 63–65^. Furthermore, knockdown of *exocyst component 3 (exoc3),* a negative regulator of pluripotency whose mammalian homolog *Tnfaip3* promotes embryonic stem cell differentiation, causes expansion of the S/G2/M (X1) neoblast fraction^66^. We speculate that these genes may be required for cell fate specification, and that their inhibition shifts neoblast dynamics in favor of renewal divisions that expand the stem cell compartment. By contrast, *apob* RNAi does not cause accumulation of S/G2/M neoblasts, but rather increases post-mitotic progeny number, and causes greater dysregulation of transcripts associated with differentiation and mature cell states. Therefore, ApoB is likely required to drive post-specification stages of differentiation, and inhibition delays late commitment steps (i.e., after cell cycle exit), and/or transitions to final mature states (Fig. 6).

Differentiation requires extensive changes in gene expression that are often preceded by genome-wide chromatin remodeling^67, 68^. Intriguingly, FA oxidation is a significant source of carbon for acetyl-CoA production and histone acetylation^69^. Consistent with a role for NLs in planarian differentiation, knockdown of *exoc3* reduces triglyceride levels, causes expansion of the neoblast population, inhibits organogenesis, and reduces expression of differentiation markers; palmitic acid supplementation rescues these differentiation-associated phenotypes^66^. Furthermore, in the sexually reproducing *S. mediterranea* planarian biotype, inhibition of *nuclear hormone receptor 1* causes NL accumulation and blocks differentiation of gonads and accessory reproductive tissues, a phenotype that is rescued by supplementation with either acetyl-CoA or Acyl-CoA synthetase^70^. Because acetyl-CoA can also enter the citric acid cycle to produce alpha-ketoglutarate, a substrate for histone demethylation^71^, ApoB inhibition could dysregulate epigenetic changes through multiple pathways. Histone acetylases, deacetylases, methyltransferases, and demethylases are conserved in planarians, and their inhibition disrupts stem cell maintenance, differentiation, and regeneration^64, 65, 72–75^. Because *apob* RNAi results in widespread dysregulation of thousands of transcripts associated with differentiating progeny, it is reasonable to suggest that in planarians, intestinal lipid stores serve as a ready carbon source that is trafficked by ApoB-containing LPs to neoblasts and progeny to support epigenetic modifications required for differentiation.

ApoB depletion may also delay differentiation by other mechanisms. For example, NL-derived fatty acids may be utilized to produce ATP via beta-oxidation and the mitochondrial electron transport chain (ETC) to support energy-dependent processes during differentiation. Consistent with this idea, planarian mitochondrial mass is higher in differentiating progeny, and pharmacological inhibition of the ETC promotes pluripotency and neoblast colony expansion, which may also limit differentiation^76^. In mammals, disrupting the ETC blocks differentiation of cardiomyocytes and mesenchymal stem cells^77, 78^, but the ETC is dispensable for differentiation of mammalian epidermal progenitor cells and *Drosophila* ovarian stem cells^79, 80^. Further study will be needed to determine whether LP-transported NLs serve as a significant energy source during planarian differentiation. Additionally, LP-mediated transport of morphogens like Hedgehog or Wnt proteins^81^, whose planarian orthologs play important roles in regulating axial polarity and tissue differentiation^53^, may be affected by *apob* knockdown. However, we find that ApoB inhibition delays, but does not block or alter regeneration of axial polarity, and we find little evidence of dysregulation of polarity-related transcripts (Supplementary Data 4 and 5), suggesting that planarian LPs may not play a major role in planarian morphogen trafficking. Similarly, fat-soluble vitamins known to influence stem cell dynamics are also transported in LPs^82–87^. Although characterization of LP cargo may yield additional insights, the dramatic dysregulation of lipid metabolism at the gene expression level, and the minimal disruption of vitamin-related gene expression in *apob(RNAi)* animals (Supplementary Data 4 and 5), suggest that LP-mediated vitamin transport may not play a significant role in planarian differentiation. Lastly, we find that *apob* inhibition dysregulated genes associated with muscle differentiation and function (Supplementary Data 5). In planarians, muscle cells not only secrete axial polarity cues, but also serve a fibroblast-like role by secreting most components of the extracellular matrix, whose functions are required to both spatially restrict the stem cell compartment, and modulate proliferation and differentiation^63, 88, 89^. *apob* RNAi causes moderate downregulation of most fibrillar collagens, as well as the basement membrane *collagen4-1,* which promotes differentiation^63^ (Supplementary Data 4 and 5). Thus, ApoB depletion may also delay differentiation indirectly, by compromising the generation and/or function of muscle cells.

The effect of *apoB* knockdown on regeneration suggests several future directions. First, although the existence of prominent LDs in planarian neoblasts has been known for decades^90^, the roles of this intriguing organelle have not been investigated. We did not assess neoblast LD numbers or size, but *apoB* knockdown dramatically reduces NL content in both X1 (S/G2/M) and X2 (G0/G1) neoblast fractions, suggesting that ApoB and LPs may influence neoblast LD content and/or function. In addition, NL content is lower in G0/G1 cells, suggesting that a primary role of LDs may be to support differentiation. Once thought of as static storage particles, recent work has demonstrated that LDs in animal cells are dynamic, multi-functional organelles that regulate nutrient sensing, cell stress responses, and even intracellular localization of histones, transcription factors, and other proteins^2, 10^. Additionally, although emerging data suggest LDs as a potential therapeutic target in cancer stem cells ^91, 92^, our knowledge of the regulation and functions of LDs in stem cells, especially during regeneration, is limited. Studies in planarians and other regeneration models could further illuminate the roles of this organelle. Second, coordinated metabolic shifts between glycolysis and oxidative phosphorylation may be a widespread aspect of stem cell transitions between quiescence, proliferation, and differentiation, and extrinsic lipids can influence these states^11–14^. Our RNA-Seq data suggest that transcription of regulators of amino acid metabolism, glycolysis, the tricarboxylic acid cycle, and other metabolic pathways respond to *apob(RNAi)*, suggesting that planarian stem cell metabolism is just as dynamic as in other animals. The fact that *apob(RNAi)* seems to primarily affect post-mitotic states also suggests that planarian neoblasts might rely primarily on glycolysis for energy and metabolite supply, and shift to lipid metabolism and oxidative phosphorylation during differentiation, as in other systems^93–95^. Again, studies in animals with high regenerative capacity could generate greater insights into whether and how injury can induce metabolic switching. Third, we find that expression of *apob-1* and *apob-2,* as well as numerous other regulators of NL metabolism are dynamically up- and down-regulated at the transcript level during regeneration (Supplementary Fig. 3E). This suggests that coordination of lipid metabolism is part of a genome-encoded program of regenerative gene expression. Unraveling which transcription factors, chromatin modifiers, and other regulators are responsible for such regulation is thus a third priority for future study.

Finally, because *apob-1* and *apob-2* are downregulated by inhibition of *nkx2.2,* an intestine-enriched transcription factor also required for regeneration^39^, our results provide a specific example of how the planarian intestine can non-autonomously influence neoblast dynamics. Intriguingly, unlike *nkx2.2* RNAi, *apob* knockdown does not reduce the abundance of phosphoHistone-H3-S10-positive neoblasts, indicating that the proliferative defect in *nkx2.2(RNAi)* animals is not caused by *apob* reduction, and additional downstream regulators of proliferation remain to be discovered. Intriguingly, dozens of additional regulators of lipid metabolism and transport of other metabolites are downregulated in *nkx2.2(RNAi)* animals (Supplementary Data 1b), suggesting additional ways in which the intestine could influence neoblasts and their niche.

In summary, we have identified *apolipoprotein B* orthologs and neutral lipid metabolism as important regulators of stem cell differentiation and regenerative tissue growth. Since the discovery of lipoproteins a century ago, their roles in lipid transport and disease have been extensively investigated^96^, but functions in stem cell regulation are not nearly as well characterized^97–100^. Efforts to define functions of LPs and their metabolic derivatives more precisely in planarians and other models will therefore improve our understanding of metabolic requirements of stem cell-driven regeneration. In addition, because lipid metabolism is amenable to pharmacological manipulation^101^, further study may provide new insights relevant to the dual goals of promoting repair of damaged human tissues, and inhibiting growth in pathological contexts like cancer.

## Methods

### Ethics Statement

Anti-ApoB-1 antibodies were generated by GenScript USA (Piscataway, NJ), an OLAW, AAALAC, and PHS-approved vendor. GenScript’s animal welfare protocols were approved by OMRF IACUC (17-58). No other vertebrate organisms were used in this study.

### Planarian care

Asexual *Schmidtea mediterranea* (clonal line CIW4)^102^ were maintained in 0.5 g/L Instant Ocean salts with 0.0167 g/L sodium bicarbonate dissolved in Type I water^103^, and fed with beef liver paste. Planarians were starved 7-10 days prior to initiating RNAi. Animals were 2-5 mm in length for most experiments except flow cytometry, for which 5-10 mm animals were used. Uninjured, intact animals were randomly selected from large (300-500 animal) pools.

### Cloning and expressed sequence tags

Transcripts were cloned as previously described^104^. Sequences were identified in the dd_Smed_v6 transcriptome^105^ and the Smed_ESTs3 library^106^. These included *nkx2.2* (dd_2716_0_1/PL08007A2A07), *apob-1* (dd_636_0_1/PL06004B2E09), *apob-2* (dd_194_0_1/PL08004B1B10), *ldlr-1* (dd_9829_0_1), *ldlr-2* (dd_5596_0_1/PL04021A1C10), *vldlr-1* (dd_1510_0_1/PL05007B1H03), *notum* (dd_24180_0_1), *wnt11-2* (dd_16209_0_1), *choline acetyltransferase/ChAT* (dd_6208_0_1), *laminin* (dd_8356_0_1/PL030015A20A02), *cathepsin La/ctsla* (dd_2567_0_1/PL06020B2D09), *piwi-1* (dd_659_0_1/PL06008A2C06), *tgs-1* (dd_10988_0_1), *solute carrier family 22 member 6/slc22a6* (dd_1159_0_1), and Niemann-Pick type C-2/*npc2* (dd_73_0_1/PL030001B20C07). *S. mediterranea ldlr-1, ldlr-2,* and *vldlr-1* were identified by BLAST homology and named after their top human refseq_protein BLASTX hits. Sequences of primers and expressed sequence tags are available in Supplementary Data 6.

### Domain organization and phylogenetic analysis

For ApoB-1, ApoB-2, Ldlr-1, Ldlr-2, and Vldrl-1, protein domains were identified using HMMSCAN (https://www.ebi.ac.uk/Tools/hmmer/search/hmmscan) to search Pfam, TIGRFAM, and Superfamily databases, with Phobius for transmembrane and signal peptide predictions (conditional E-value cutoff of 1e-03)^107^. For the ApoB phylogenetic tree, N-terminal Vitellogenin domains from ApoB and related proteins were aligned in Geneious using MAAFT (default settings), and alignment was manually trimmed to the N- and C-terminal boundaries of human Apo B-100. Phylogenetic tree was generated using PhyML 3.0^108^ (http://www.atgc-montpellier.fr/phyml/), using AIC for automatic selection of the LG substitution model, BioNJ starting tree, NNI for tree topology improvement, and 100 bootstrap replicates. Accession numbers for proteins used in domain diagrams and phylogenetic analysis are included in Supplementary Data 6.

### In situ hybridization

Riboprobe synthesis, WISH, and FISH were conducted as previously described^104^. Riboprobes were used at 0.05-1 ng/μl. Cryosections were generated after FISH as in^109^.

### RNAi

dsRNA synthesis and RNAi experiments were conducted as described^41, 104^ by mixing one μg of in vitro-synthesized dsRNA with 8-9 μl of 1:10 food coloring:water mix, and 40 μl of 2:1 liver:water homogenate. For *nkx2.2* RNAi, animals were fed only once; RNA was extracted for RNA-Seq after seven days. For *apob* RNAi, 2 μg control *egfp* dsRNA or 1 μg each *apob-1* and *apob-2* (for simultaneous RNAi) were mixed with liver and food coloring, and animals were fed 5 times for initial viability experiments, and 3-5 times for most other experiments, separating animals with “mild” (*apob-M)* and “severe” *(apob-S)* phenotypes prior to fixation, amputation, or flow cytometry. Non-eating planarians were always removed from the experiment if they refused a second dsRNA feeding one day later.

### pH3-PS10 immunolabeling

Mucus removal and fixation were conducted with 2% ice-cold HCl (3 min) and methacarn at room temperature (RT, 20 min), followed by bleaching in 6% H2O2 in methanol as in^109^. Fixed animals/regenerates were blocked (4 hr, RT) in IF block (1X PBS, 0.45% fish gelatin, 0.6% IgG-free BSA, 0.3% Triton X-100), incubated in rabbit anti-phospho-Histone H3-S10 at 1:2000 overnight (O/N, 4°C) (Cell Signaling 3377S), washed 8X in PBSTx (1X PBS plus 0.3% Triton X-100) (30 min each, RT), re-blocked for 1 hr, then incubated with goat anti-rabbit-HRP (1:2000) (Jackson ImmunoResearch 111-035-003) (O/N, 4°C). Samples were again washed 8X (20-30 min each, RT), then tyramide signal amplification (TSA) was conducted for 10 min as described^110^ with TAMRA-tyramide. Samples were washed in PBSTx for two days, then mounted in Vectashield (Vector Labs H-1000-10).

### Immunolabeling (cryosections)

Cryosections (12 μm) of planarians relaxed in 0.66M MgCl2 and fixed (O/N, 4°C) in 4% formaldehyde (EM)/1X PBS were generated as described^109^. After rehydration, heat-mediated antigen retrieval (10 min) in 10 mM sodium citrate, pH 6.0 was performed. Slides were permeabilized for 30 min in 1X PBS/0.2% Tween-20, then blocked for 30 min at RT with 0.45% fish gelatin and 0.6% BSA for 30 min in PBSTw (1X PBS, 0.05% Tween 20). Slides were incubated with custom rabbit anti-ApoB-1 (1:1000, 0.59 μg/ml) and mouse 6G10 anti-muscle (1:250 in block)^111^ in blocking buffer (O/N, 4°C). Slides were washed three times (10 min, RT) with PBSTw after antibody incubation. Slides were then incubated with goat anti-rabbit-HRP (1:2000) (Jackson ImmunoResearch, 111-035-144) and goat anti-mouse-488 (1:250) (Jackson ImmunoResearch, 115-545-146) at RT for 60 min. Slides were washed three times (in PBSTw), with DAPI (2 μg/ml) counterstaining during the first wash. TSA was conducted for 10 min with TAMRA-tyramide as described^110^, followed by washes. Slides were mounted in Fluoromount G (Southern Biotech).

### Anti-ApoB-1 antibody generation

Sequence corresponding to the N-terminal Vitellogenin domain of ApoB-1 (dd_636, nucleotides 45-1919, amino acids 1-625) was supplied to GenScript USA (Piscataway, NJ). GenScript synthesized a plasmid encoding this region fused to KLH and a HIS tag, expressed and purified the fusion protein, immunized New Zealand rabbits, affinity purified the antibody, and assessed antibody titer and immunogenicity with both ELISA and Western blot.

### Protein extraction and Simple Western analysis

5-10 planarians were rocked gently (40-50 rpm) for 7 min in 7.5% N-Acetyl-L-Cysteine and rinsed 2X in 1X PBS. Samples were then homogenized using a motorized Kontes pestle grinder in 250 μl RIPA (50 mM Tris pH 8.0, 150 mM NaCl, 1% NP-40, 0.5% sodium deoxycholate, 0.5% SDS) with 40 mM DTT and 1X Halt Protease Inhibitor cocktail (Thermo Scientific 78430). After 30 min on ice, samples were centrifuged (20,817 x *g,* 15 min, 4°C), and supernatant was recovered and stored at -80°C. DTT concentration was reduced by buffer exchange with RIPA (1 mM DTT) using Amicon Ultra 3 kDa columns (UFC500396), then protein concentration was determined using a BCA kit (Pierce 23225) and DeNovix cuvet spectrophotometer according to manufacturers’ protocols. For Simple Western (ProteinSimple), lysates were run according to the manufacturer’s protocol on a Wes Instrument (running Compass v4.0.0) at 0.1 mg/ml using the 66-440 kDa Wes Separation module (SM-W008) with anti-ApoB-1 at 1:250 (∼2.3 μg/ml). ApoB-1 peak areas were calculated with the following settings: Range 10-600; Baseline Threshold 1.0, Window 15.0, and Stiffness 1.0; and Peak Find Threshold 10.0 and Width 20.0 using the “Dropped Lines” method. Settings were identical for total protein peak area, with these exceptions: Baseline Window 50.0 and Stiffness 0.5. For the regeneration time course, ApoB-1 protein levels were run in biological triplicates, normalized to total protein using the Wes Total Protein Detection Module (DM-TP01), and then normalized to the “0 hour” time point in Excel.

### Oil Red O staining

Planarians were relaxed in 0.66 M MgCl2, fixed overnight (RT) in 4% formaldehyde (EM grade) in 1X PBS, protected in sucrose, and cryosectioned (20 μm) onto SuperFrost Plus slides^109^. Slides were rehydrated in deionized (DI) water (3 x 10 min, RT), then stained in Oil Red O (Sigma O0625) solution (6 ml Whatman-filtered 0.5% Oil Red O in 100% isopropanol plus 4 ml ultrapure water) for 15 min at RT. Slides were quickly dipped 3-5X in 60% isopropanol to remove excess dye, then rinsed for 1 min in 60% isopropanol, then rinsed 1 min in DI water. Slides were then rinsed in PBS-Tween-20 (0.01%, to prevent drying), then mounted in 90% glycerol/1X PBS, and imaged within 2-3 days.

### Thin Layer Chromatography

Ten planarians (5-8 mm) were placed in 1.7 ml microcentrifuge tubes with all planarian salts removed, and the animals’ mass was obtained. Lipids were then extracted using the Folch method^112^. Briefly, 1 ml ice-cold 2:1 chloroform:methanol was added, then animals were sonicated in ice water in a cup-horn sonicator (10 cycles of 5 second pulses at ∼48-55W). Samples were rocked at RT for 5 hr, then centrifuged (2 min at 16,000 x *g,* 4°C) to pellet insoluble material. Supernatant was recovered to a new tube and stored at -80°C. 1 ml 2:1 chloroform:methanol was added to the pellet, re-sonicated, rocked overnight at RT, then centrifuged as above. 0.2 volumes 0.9% NaCl (in water) was added to each extract, tubes were inverted gently 10-15X to mix, vortexed for 10-15 sec, then centrifuged (2 min at 2,000 x *g,* RT). Lower phases from each biological replicate (4 control and 4 *apob(RNAi)* replicates, 10 animals each) were recovered, and speed-vacuumed (30°C x 60-90 min with spinning) to evaporate solvent. Concentrated lipids were resuspended in 2 μl chloroform per mg animal mass (above) and stored at -80°C (for less than 7 days). TLC was performed as previously described^113^ with slight modifications. 150Å silica gel HL 250 μl 20x20 cm plates (iChromatography 66011/Miles Scientific P76011) were pre-equilibrated with 1:1 chloroform:methanol (∼ 1 hr). After drying, 1 μl lipids and 3 μl standards (30 μg mono-, di-, triglyceride mix, SUPELCO 1787-1AMP, plus 30 μg cholesteryl palmitate, SIGMA C6072) were spotted onto the plates using TLC spotting capillaries. Non-polar lipids were resolved with a 80:10:1 petroleum ether:ethyl ether:acetic acid mix. After drying, TLC plates were sprayed with Primuline (SIGMA 206865) (1 mg/ml fresh stock in ultrapure water, diluted 1:100 into 80 ml acetone plus 20 ml ultrapure water), dried, and imaged on an Alpha Innotech chemiluminescent imager with Cy2 excitation/emission filters. Peak areas in images were quantified in ImageJ/Fiji using the “Plot Lanes” function in the Gels submenu (https://imagej.nih.gov/ij/docs/menus/analyze.html). Averages were calculated and normalized to controls in Excel.

### RNA extraction, library preparation, and RNA sequencing

Uninjured planarians and regenerating tissue fragments were homogenized in Trizol using a motorized Kontes pestle grinder, and RNA was extracted using two chloroform extractions and high-salt precipitation buffer according to the manufacturer’s instructions. After precipitation, solutions were transferred to Zymo RNA columns for DNAse treatment and purification, according to manufacturer’s instructions. RNA samples were analyzed using Agilent RNA ScreenTape on an Agilent TapeStation 2200 according to manufacturer’s protocol.

For analysis of gene expression in control vs. *nkx2.2(RNAi),* mRNA was enriched using oligo-dT homopolymer beads, and libraries were generated using the Illumina Truseq Stranded mRNA Library Prep Kit according to the manufacturer’s protocol. Final libraries were assayed on the Agilent TapeStation for appropriate size and quantity. Libraries were pooled in equimolar amounts as ascertained by fluorometric analysis, then final pools were absolutely quantified using qPCR on a Roche LightCycler 480 with Kapa Biosystems Illumina Library Quantification Reagents. Paired-end (2x150 bp) sequence was generated on an Illumina NovaSeq 6000 instrument. 28M-43M reads were generated for each of three biological replicates per condition. For analysis of gene expression in control, *apob-M,* and *apob-S* animals, total RNA for 4-6 biological replicates per condition was submitted to GENEWIZ (South Plainfield, NJ) for library generation using NEB NEXT ULTRA library prep and RNA sequencing with standard Illumina adapters. Paired-end (2x150 bp) sequence was generated on an Illumina HiSeq 4000 instrument; 19M - 26M reads were generated for each replicate.

### Read mapping

For both *nkx2.2(RNAi)* and *apob(RNAi)* experiments, quality control and read mapping to unique transcripts in dd_Smed_v6^104, 105^ were conducted with FastQC (v0.11.5)^114^, BBDuk (v35.66) (https://sourceforge.net/projects/bbmap/), and Bowtie2 (v2.3.1)^115^. BBDuk (v36.99) settings for paired end reads: k=13 ktrim=r mink=11 qtrim=rl trimq=10 minlength=35 tbo tpe. Bowtie2 (v2.3.1) for paired end reads was used for mapping, with “-a” multi-mapping and “--local” soft-clipping allowed. For read summarization, the “featureCounts” utility in the Subread package (v1.6.3)^116^ was used with a custom “.SAF” file and options “-p -M -O -F SAF” to include multi-mapping and multi-overlapping reads.

For mapping of X1/X2/Xins and PIWI-HI/-LO/-NEG bulk sequence, regeneration fragment sequence, and whole animal 24 hr post-irradiation sequence^49^, fastq files were downloaded from NCBI GEO (GSE107874), mapped to dd_Smed_v6_unique using BBDuk and Bowtie2, followed by count summarization using Samtools as previously described^104^.

### Differential expression analysis

Read counts matrices were imported into R, then analyzed in edgeR v3.8.6^117^. First, all transcripts with counts per million (CPM) < 1 in three samples (*nxk2.2(RNAi),* Zeng X1/X2/Xins data, and Zeng 24 hr irradiation data) or four samples (all others) (e.g., lowly expressed transcripts) were excluded from further analysis. Next, after recalculation of library sizes, samples were normalized using trimmed mean of M-values (TMM) method, followed by calculation of common, trended, and tagwise dispersions. Finally, differentially expressed transcripts were identified using the pairwise exact test (*nkx2.2(RNAi)* and *apob(RNAi)* experiments, Zeng 24 hr irradiation data) or the generalized linear model (GLM) likelihood ratio test (other Zeng datasets). Expression changes were considered to be significant if the false discovery rate adjusted *p* value (“FDR”) was <.05.

### Gene Ontology analysis

GO analysis was conducted using BiNGO^118^ using a custom *S. mediterranea* GO annotation as previously described^104^. RefSeq protein collections used for BLASTX and UniProtKB Biological Process GO terms used for annotation were downloaded for each organism in April 2020.

### Hierarchical clustering and heat maps

Hierarchical clustering of transcripts annotated with lipid-metabolism-related GO terms was conducted using EdgeR-generated log2FC values in Cluster 3.0^119^, with Euclidean distance and complete linkage. Heat maps were generated with Java Treeview^120^.

### qRT-PCR

Total RNA was extracted from biological triplicates (5-10 animals/fragments per replicate) using Trizol as for RNA-Seq samples. RNA was reverse transcribed using the iScript cDNA Synthesis kit (BioRad 1708890). *apob-1* and *apob-2* levels were detected using the Fast Start Essential Green DNA master mix (Roche 06924204001) on a Roche LightCycler 96 instrument. RNA levels were normalized to the geometric mean of endogenous controls *ef-2* and *gapdh* using the Livak ΔΔCt method^121^.

### Flow cytometry

Planarians were dissociated and filtered in CMFB with collagenase as described^62^. Cells were labeled at RT with Hoechst 33342 (50 μg/ml) for 45 min, followed by addition of propidium iodide (1 μg/ml). For neutral lipid labeling, BODIPY 493/503 (Molecular Probes D3922) at 10 ng/ml was included with Hoechst. Cells were analyzed on a Becton Dickinson FACSCelesta instrument with 405 nm, 488 nm, and 640 nm lasers. After gating for live cell singlets (Supplementary Fig. 9a-c), X1, X2, and Xins gates were drawn using two criteria: cell proportions were approximately 15% (X1) 25% (X2) and 60% (Xins), and reductions in X1 and X2 fractions in 4-day post-irradiation animals were >95% and ∼70%, respectively. Data were analyzed and plots were generated in FloJo (v10.7.1). For irradiation, uninjured planarians were dosed with 60 Grays (6,000 rads) using RS-2000 Biological Research X-Ray Irradiator (Rad Source, Buford, GA).

### Cross-referencing of *apob(RNAi)* RNA-Seq data with published transcriptomes

For comparison of dysregulated transcripts in *apob-M* and *apob-S* animals with bulk neoblast transcriptome data^49^, we first identified “signature” transcripts as follows. “X1 signature” transcripts were defined as those with a log2FC>0 (FDR<.05) compared to both “X2” and “Xins.” “X2 signature” transcripts had log2FC>0 (fdr<.05) vs. both “X1” and “Xins.” “Xins signature” had log2FC>0 (fdr<.05) vs. both “X1” and “X2.” Similarly, “PIWI-HI signature” transcripts were defined as those with a log2FC>0 (fdr<.05) compared to both “PIWI-LO” and “PIWI-NEG.” “PIWI-LO signature” transcripts had log2FC>0 (fdr<.05) vs. both “PIWI-HI” and “PIWI-NEG.” “PIWI-NEG signature” transcripts had log2FC>0 (fdr<.05) vs. both “PIWI-HI” and “PIWI-LO.” Next, we used the “merge” function and “VennDiagram” package in RStudio (v1.2.1335) to identify signature transcripts in X1/X2/Xins or PIWI-HI/-LO/-NEG that were also dsyregulated (up or down) in *apob-M* or *apob-S* animals, as shown in Fig. S6. “% of dysregulated transcripts” was calculated as the number of overlapping transcripts divided by the total number of X1/X2/Xins or PIWI-HI/-LO/-NEG transcripts.

For comparison of dysregulated transcripts in *apob-M* and *apob-S* animals with single cell type/state data in^40^, we again used the “merge” function in RStudio to identify the number of transcripts enriched in individual lineage subclusters (Table S2)^40^ that were also dysregulated by *apob* RNAi. “% of dysregulated transcripts” was calculated as the number of overlapping transcripts divided by the total number of transcripts in each individual subcluster. In Fig. 5, the “N/TS” (<u>N</u>eoblast/<u>T</u>ransition <u>S</u>tate) designation included subclusters with high *piwi-1* mRNA expression thought to be neoblast/progenitor subpopulations in epidermal, intestine, and protonephridia lineages based on the conclusions of Fincher et al. and other published data^45, 62, 122–124^. Similarly, the “P” (<u>P</u>rogeny) and “M” (<u>M</u>ature) designations were based on conclusions from both single cell RNA-Seq data and previous work. For lineages that are less well understood *in vivo* (Fig. S8), we designated subclusters/states using both transcript dysregulation in 24 hr irradiated animals ^49^ and *piwi-1* mRNA levels in t-SNE plots^40^. “N/TS” subclusters possessed the greatest number of irradiation-dysregulated transcripts and the highest *piwi-1* expression; “P” subclusters possessed fewer (by proportion) irradation-dysregulated transcripts and lower *piwi-1* expression; and “M” subclusters had the fewest radiation-sensitive transcripts and negligible *piwi-1* expression.

### Image Collection and Quantification

Epifluorescent images (FISH samples) were collected on a Zeiss AxioObserver.Z1 with Excelitas X-Cite 120 LED Boost illumination and Zen 2.3 pro. For quantification of pharynx and brain size, organ area and animal area were measured in ImageJ^125^ and organ-to-body size ratios were calculated. For intestine, length of anterior branch and posterior branches were measured in ImageJ. Means of posterior branch length were calculated, and then posterior-to-anterior (head fragments) or anterior-to-posterior (tail fragments) length ratio was calculated. For tail fragments with split anterior branch, anterior branch length was measured from the anterior of the pharynx to the tip of the anterior-most primary branch. For tail fragments, the split anterior branch phenotype (failure to fuse at the midline) was scored if there was an obvious gap between anterior branches for at least half the length of the anterior branch. For images of *apob-1* and *apob-2* FISH on cryosections, z-stacks were collected with an Apotome.2 for generation of maximum orthogonal projections.

For anti-pH3-PS10-labeled samples, z-stacks were collected on a Zeiss AxioObserver.Z1 at 10X magnification, followed by tile stitching, extended depth of focus projection, and background subtraction (PS10 channel only) with a radius of 30. Control and experimental samples to be compared were imaged at identical exposures. For quantification, animal area and PS10+ nuclei were quantified using the Automated Segmentation tools in Zen 2.3 pro. Briefly, animal area was measured with Gaussian smoothing, no background subtraction or sharpening, Morphology separation, and custom threshold settings that were the same for all samples to be directly compared. PS10+ nuclei number was measured using Lowpass smoothing, Rolling Ball background subtraction, Delineate sharpening, Watersheds separation, with threshold settings and other parameters that were identical for all samples to be directly compared. Number of mitoses per area were calculated for each animal/fragment.

Confocal images were collected on a Zeiss LSM 710 or LSM 880 laser scanning microscope with 10X Plan NeoFluar, 20X Plan Apo, or 40X C-Apo objectives. For orthogonal projections (anti-ApoB1 immunolabeling and *ldlr/vldlr* FISH), between three and ten z-planes were collected at 1-2X “optimal” section thickness (based on objective NA). For *notum* and *wnt11-2* FISH, full fragment thickness stacks were projected to ensure that all mRNA-positive cells were counted.

Images of Oil-Red-O-stained sections, live animals, live regenerates, and WISH samples were collected on a Zeiss Stemi 508 with an Axiocam 105 color camera, or a Zeiss Axio Zoom.V16 with an Axiocam 105 camera. In some cases brightness and/or contrast were adjusted in Adobe Photoshop to improve signal contrast.

### Statistics

Detailed data and information regarding statistical testing are included in Source Data 2. For experiments with statistical analysis, n values are indicated exactly by the number of data points in figure plots, along with definitions of error bars and *p* or *q* values; all tests were conducted in Prism 9 (GraphPad Software, San Diego, CA). For one-way ANOVA, ordinary ANOVA was performed unless Brown-Forsythe and Bartlett’s tests indicated standard deviations were significantly different, then Brown-Forsythe and Welch ANOVA tests were performed. For two-way ANOVA (flow cytometry experiments with irradiation), *q* values (FDR-adjusted *p* values) were reported when interaction between RNAi condition (“Genotype”) and irradiation was significant (X2 subpopulation). Otherwise, *p* values were derived using one-way ANOVA. *p* values of <.05 (*), <.01 (**), <.001 (***), and <0.0001 (****) were annotated with asterisks in figures.

Replicate information for other experiments: polarity marker analysis, 5-15 fragments per condition; for WISH and FISH, images are representative of 3-6 individual animals or fragments; for anti-ApoB1 and Oil Red O labeling, images are representative of sections from 2-3 fragments/animals, and of at least two repeated experiments.

For statistical testing of overlap between genes dysregulated in *apob(RNAi)* planarians and other RNA-Seq data sets, the R Package GeneOverlap^126^ was used to conduct Fisher’s exact test on each comparison. Total number of detected transcripts (“genome size” in GeneOverlap) was determined conservatively by only including transcripts detected in both *apob* RNA-Seq and bulk^49^ or single cell^40^ sequencing data. For sc-RNA-Seq, the digital expression matrix in GEO GSE111764 was normalized in Seurat as in Fincher et al.^40^; only transcripts with non-zero expression in 0.5% of cells (Fincher TableS1) in each subcluster were considered to be detected. *p* values <0.05 are indicated with a caret (^) in figures.

Replicate and statistical information for RNA-Seq and other bioinformatics experiments are detailed in appropriate Methods sections.

### Data availability

Raw and processed RNA-Seq data associated with this study will be made available in the NCBI Gene Expression Omnibus (GEO) upon publication. Other data supporting this study’s findings are available within the article and its Supplementary files, or are available from the authors upon reasonable request.

### Reporting summary

Further information on research design is available in the Nature Research Reporting Summary linked to this article.

## Competing interests

The authors declare no competing financial interests.

## Acknowledgments

We are very grateful to Phil Newmark (HHMI and the Morgridge Institute for Research), in whose laboratory this project was initiated. We thank members of the Newmark and Forsthoefel labs, and OMRF colleagues Pat Gaffney, Linda Thompson, Dean Dawson, and Hui-Ying Lim for insightful discussions. We thank Jochen Rink, James Cleland, and Hanh Vu (Max Planck Institute for Biophysical Chemistry) for sharing planarian protein extraction protocols, Vasileios Morkotinis for *tgs-1* plasmid, and Rachel Roberts-Galbraith (Univ. of Georgia) for *notum* and *wnt11-2* plasmids. We thank Steve Farber (Carnegie Institution for Science) for advice on lipid extraction and TLC, and Jayhun Lee (Morgridge) for collaborative development of planarian lipid analysis protocols. We are grateful to Hadi Maktabi (ProteinSimple) and to OMRF colleagues Summer Wang and Lin Wang for help in developing Wes protocols; and to members of the OMRF Flow Cytometry Core (Jacob Bass and Diana Hamilton), the Quantitative Analysis Core (Lori Garman and Nathan Pezant), the Imaging and Histology Core, the Clinical Genomics Core, the Gnotobiotic Mouse Core (for use of X-irradiator), and IT/Research Computing Services for invaluable technical assistance. JRO was supported by the Summer Research Opportunities Program at the University of Illinois at Urbana-Champaign. This work was supported by NIH Centers of Biomedical Research Excellence (COBRE) GM103636 (Project 1 to DJF), and the Oklahoma Medical Research Foundation.

## Author Contributions

Conception and design of the project: DJF and LLW; data collection: CGB, LLW, JRO, NIC, and DJF; data analysis and visualization: CGB, LLW, NIC and DJF; data interpretation: LLW, CGB, and DJF; RNA-seq analysis: DJF; manuscript preparation: DJF, LLW, and CGB.

## Supplementary Figures and Figure Legends

**Supplementary Figure 1.**
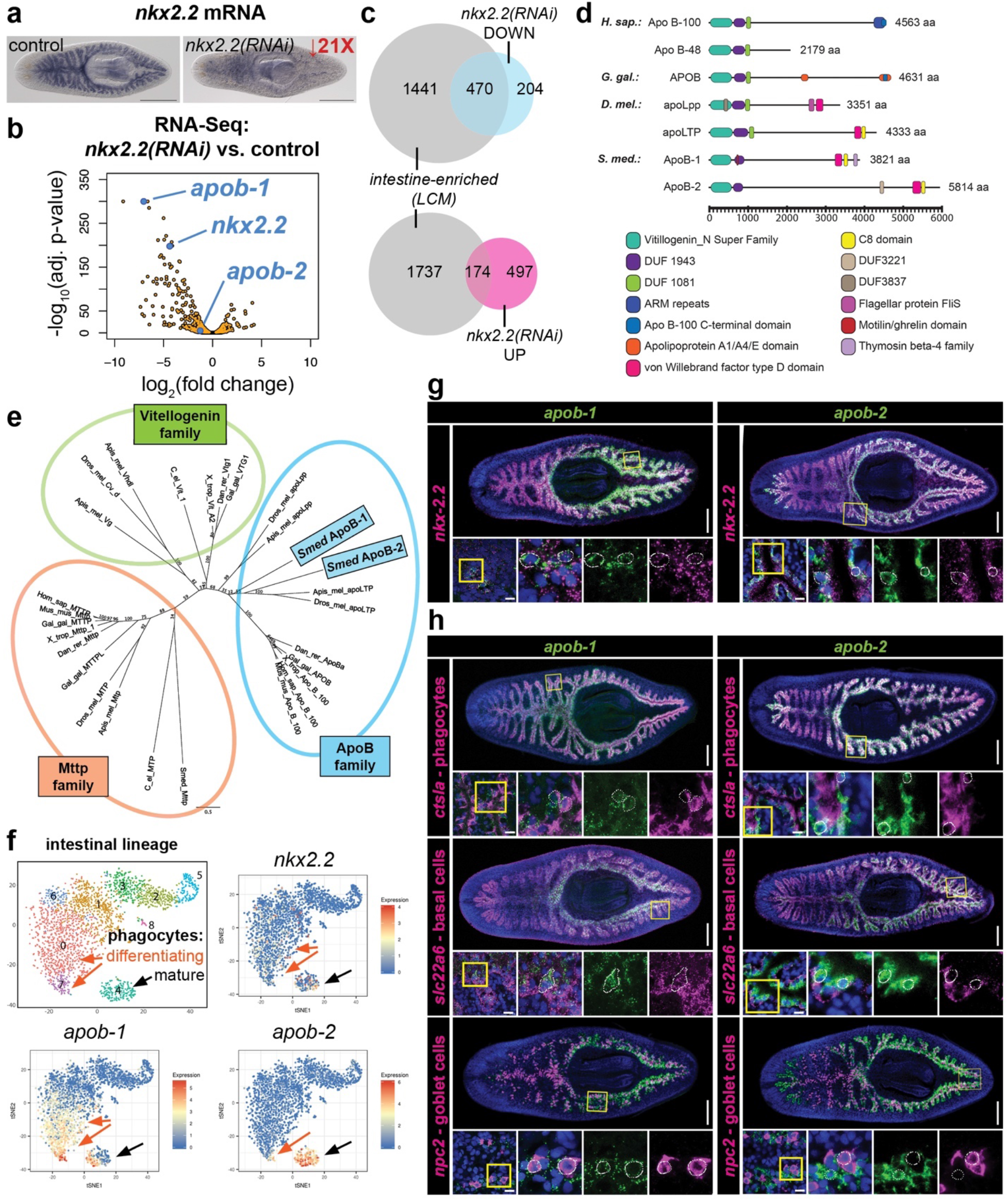
*apob-1* and *apob-2* encode intestine-enriched ApoB orthologs. **(a)** *nkx2.2* mRNA *in situ* expression patterns in uninjured control and *nkx2.2(RNAi)* planarians. Volcano plot showing downregulation of *apob-1, apob-2,* and *nkx2.2* in *nkx2.2(RNAi)* animals. An offset of 1e-300 was added to all FDR-adjusted *p* values to enable plotting of transcripts with *p*=0. **(c)** 470 downregulated (top) and 174 upregulated (bottom) transcripts in *nkx2.2(RNAi)* animals exhibited intestine enrichment in a previous study^104^. Total numbers of dysregulated transcripts in *nkx2.2(RNAi)* samples were slightly lower than in Supplementary Data 1, because some were undetectable in the intestine data set. **(d)** Conserved domains in human (*H. sap.),* chicken (*G. gal.*), fly (*D. mel.*), and planarian (*S. med.*) ApoB proteins. **(e)** Phylogenetic relationship of planarian (*Smed*) ApoB-1 and ApoB-2 (based on similarity of N-terminal Vitellogenin domains) with closely related protein families in human (*Hom_sap*), mouse (*Mus_mus*), chicken (*Gal_gal*), fly (*Dros_mel*), honeybee (*Apis_mel*), frog (*X_trop*), and *C. elegans* (*C_el*). Branch support is indicated. **(f)** t-SNE plots from single cell transcriptomes^40^ showing expression of *nkx2.2, apob-1,* and *apob-2* in the intestinal lineage. All transcripts were enriched in differentiating progeny (subclusters 0/7, orange arrows) and mature phagocytes (subcluster 4, black arrow); *nkx2.2* was also enriched in neoblasts/transition state cells (subcluster 1). **(g)** Double FISH showing co-expression of *apob-1, apob-2,* and *nkx2.2* mRNA. **(h)** Double FISH showing expression of *apob-1, apob-2* in phagocytes and basal cells (*apob-1* only), but not goblet cells. Yellow boxes indicate regions magnified in insets. Scale bars: 500 µm **(a)**; 200 µm **(g-h)**; 10 µm **(g-h** insets, bottom left).

**Supplementary Figure 2.**
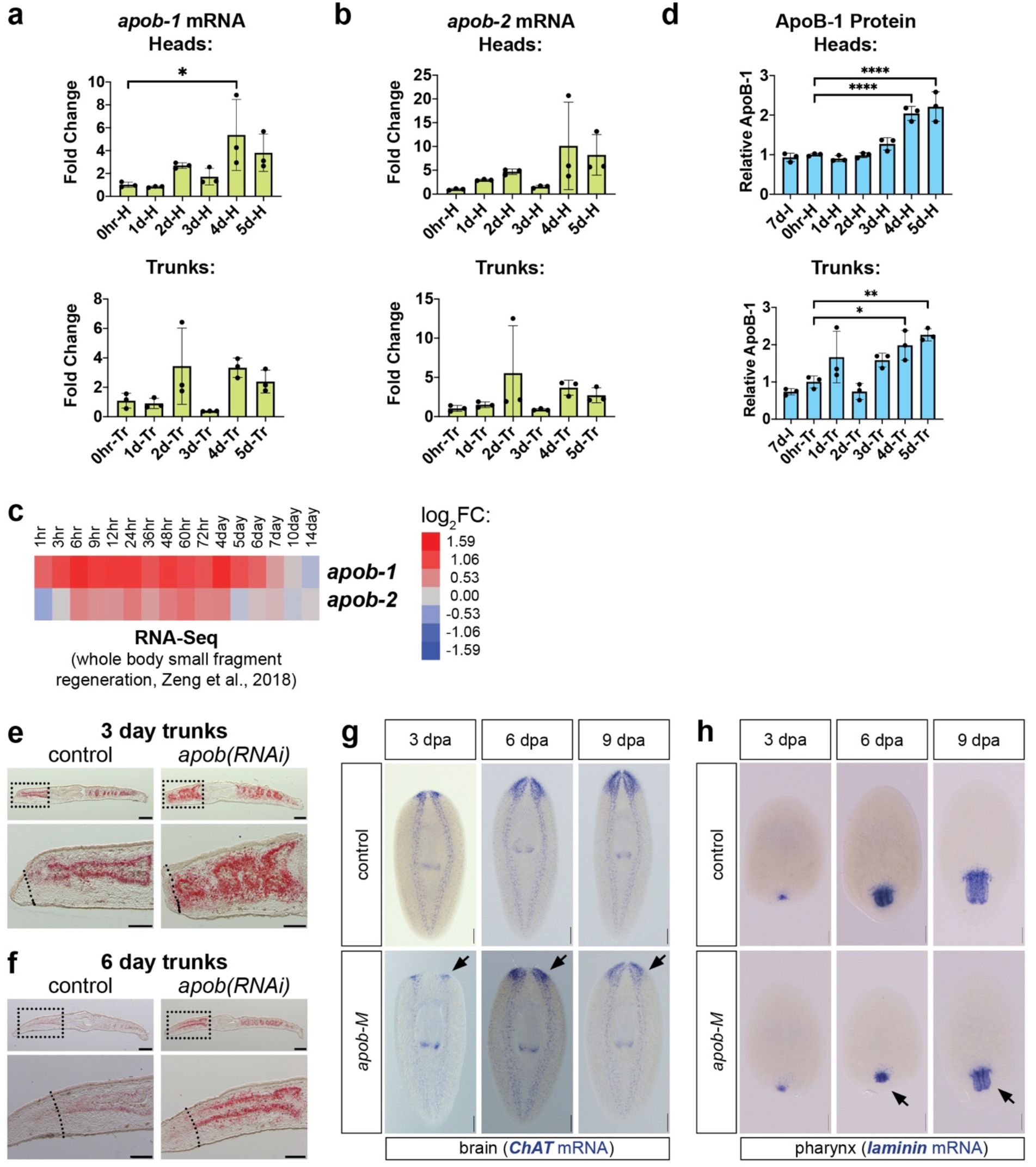
Further characterization of *apob* expression and RNAi phenotypes. **(a)** *apob-1* mRNA was upregulated in head and trunk regenerates (qRT-PCR). **(b)** *apob-2* mRNA was upregulated in head and trunk regenerates (qRT-PCR). **(c)** *apob-1* and *apob-2* mRNA levels were upregulated in whole fragment regeneration RNA-Seq data (Zeng et al., 2018). **(d)** ApoB-1 protein was upregulated during regeneration (3-5 dpa) in head and trunk fragments (Simple Western “Wes” capillary-based immunoassay). 7d-I, 7 day Intact (starved and uninjured) animals. **(e-f)** Neutral lipids accumulate in *apob*(RNAi) (“mild”) regenerates. Dashed line indicates approximate plane of amputation. Anterior is left. **(g-h)** Brain (G) and pharynx (H) regeneration were delayed, but not blocked, by *apob* RNAi. Arrows indicate smaller organs in *apob-M* animals relative to controls (representative images). Significance testing (qPCR and Wes): one-way ANOVA (comparisons to 0 hr controls) with Dunnett’s T3 multiple comparisons test. Error bars = mean +/- S.D. DAPI (blue) in **(d-g)**. Scale bars: 200 µm **(e-f,** upper panels); 100 µm **(e-f,** lower panels); 100 µm **(g-h)**.

**Supplementary Figure 3.**
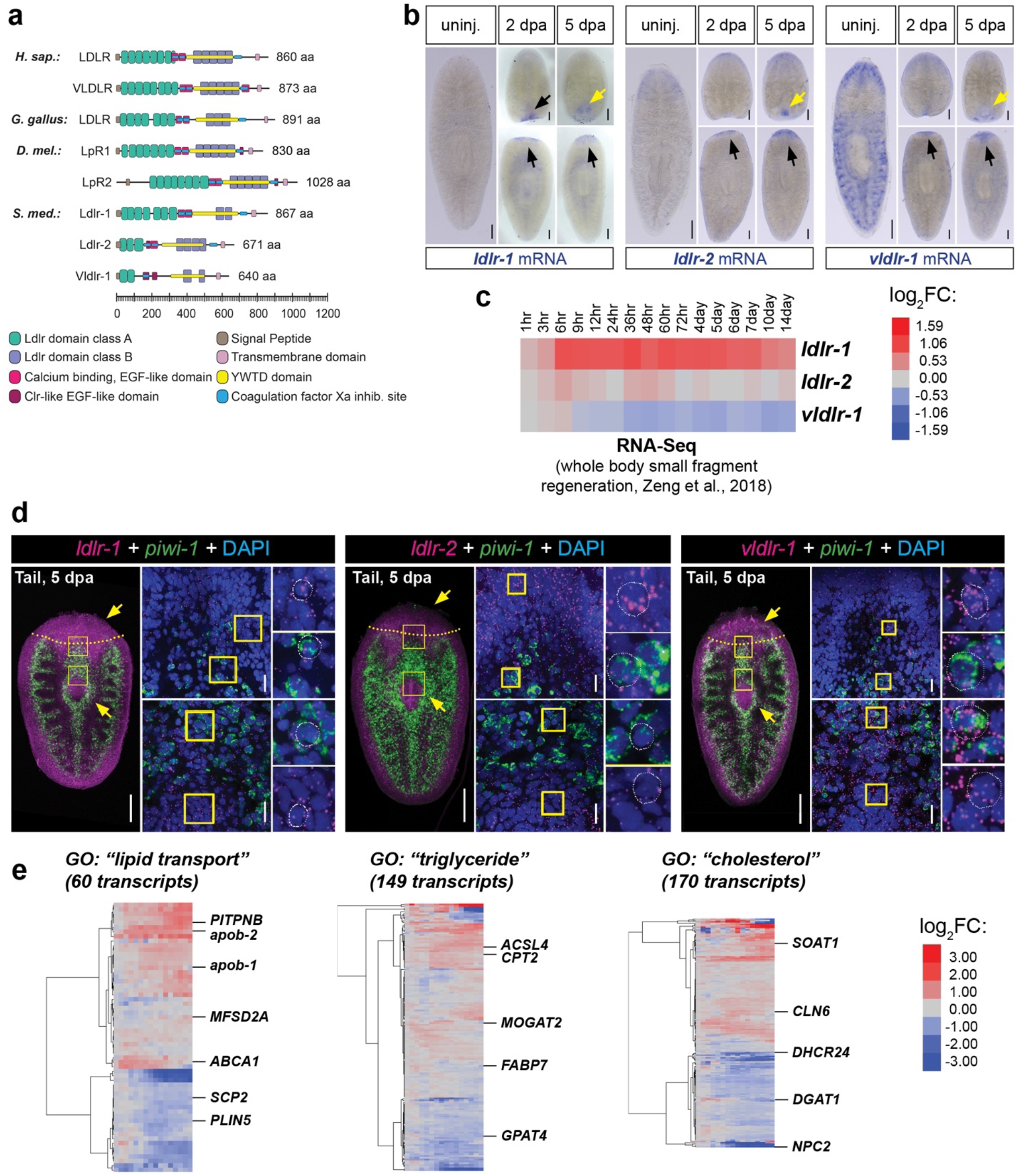
Lipoprotein receptor expression is upregulated during planarian regeneration. **(a)** Schematic of domains in LDLR and related homologs. Only domains common to Ldlr-like proteins in multiple species are shown. Clr-like EGF-like domains are shown only if they do not overlap with Ca-binding EGF-like domains. **(b)** WISH showing upregulation of planarian *ldlr* homologs in blastemas (black arrows) or developing pharynges (yellow arrows) at 2 and 5 dpa. **(c)** *ldlr-1* and *ldlr-2* mRNA levels were upregulated in whole fragment regeneration RNA-Seq data (Zeng et al., 2018). **(d)** *ldlr* homologs (magenta, mRNA FISH) were expressed in *piwi-1+* neoblasts (green, mRNA FISH) as well as differentiating (*piwi-1-*negative) cells in brain and pharynx (arrows) in 5-day regenerates. **(e)** Heat maps showing numerous planarian transcripts related to lipid metabolism that were up- and down-regulated during regeneration in whole fragment regeneration RNA-Seq data (Zeng et al., 2018). See Supplementary Data 3 for transcript IDs and expression data. Scale bars: 200 µm **(b)**; 200 µm **(d,** left panels); 20 µm **(d,** right panels).

**Supplementary Figure 4.**
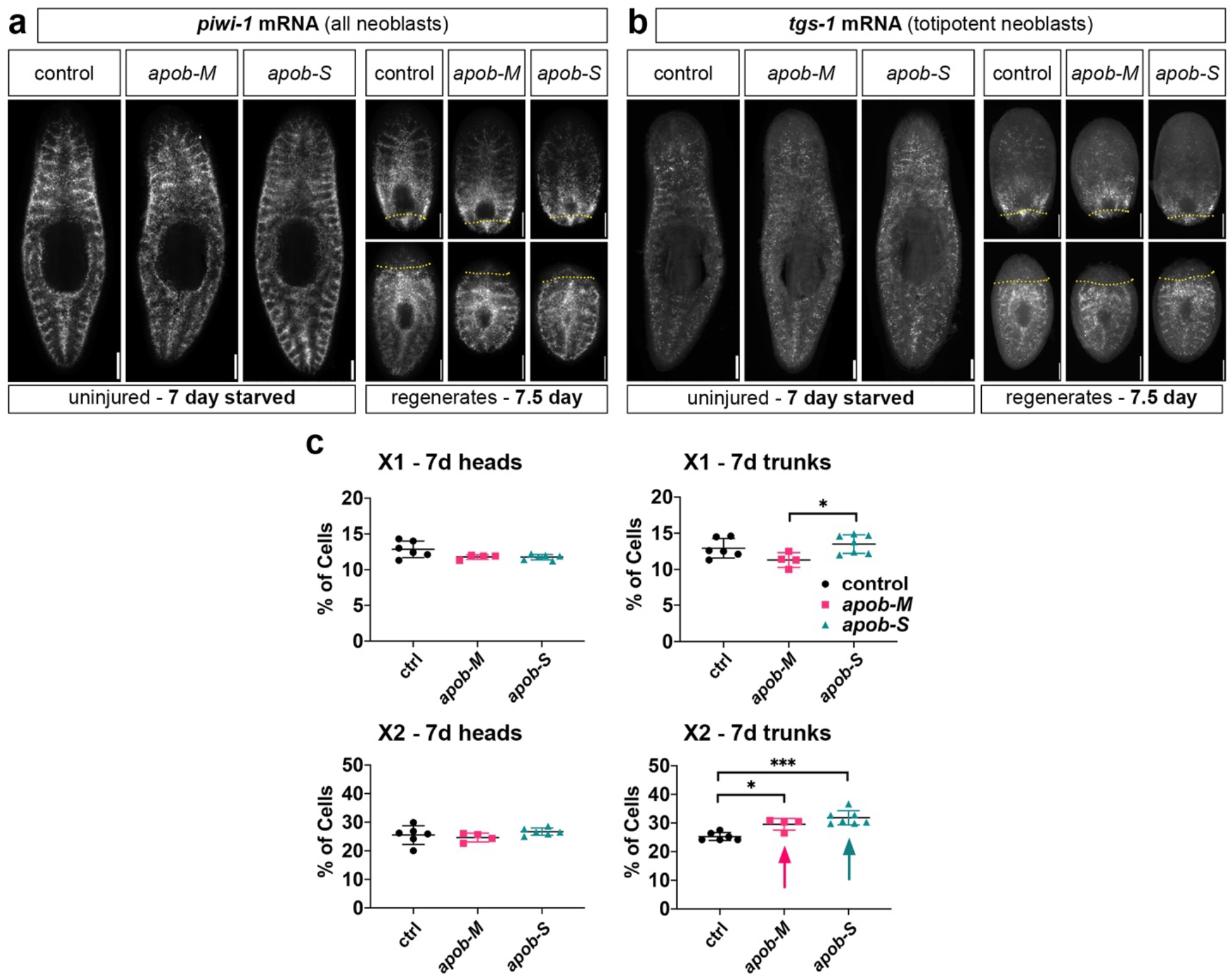
Further characterization of effects of *apob* inhibition on neoblasts and neoblast progeny. **(a)** Distribution of *piwi-1-*expressing neoblasts in uninjured (7 day starved, left panels) and 7.5 dpa head and tail regenerates (right panels). **(b)** Distribution of *tgs-1-*expressing neoblasts in uninjured (7 day starved, left panels) and 7.5 dpa head and tail regenerates (right panels). Whole-mount FISH **(a, b)**; epifluorescent images of representative samples. Dotted lines, approximate amputation plane. **(c)** Percentage of cells in X1 and X2 in 7 day head (left) and trunk (right) regenerates. Arrows indicate significant increases in X2. One-way ANOVA and Dunnett’s (X1, 7 day heads) or Tukey’s (all others) multiple comparisons test. Error bars = mean +/- S.D. Scale bars: 200 µm **(a-b)**.

**Supplementary Figure 5.**
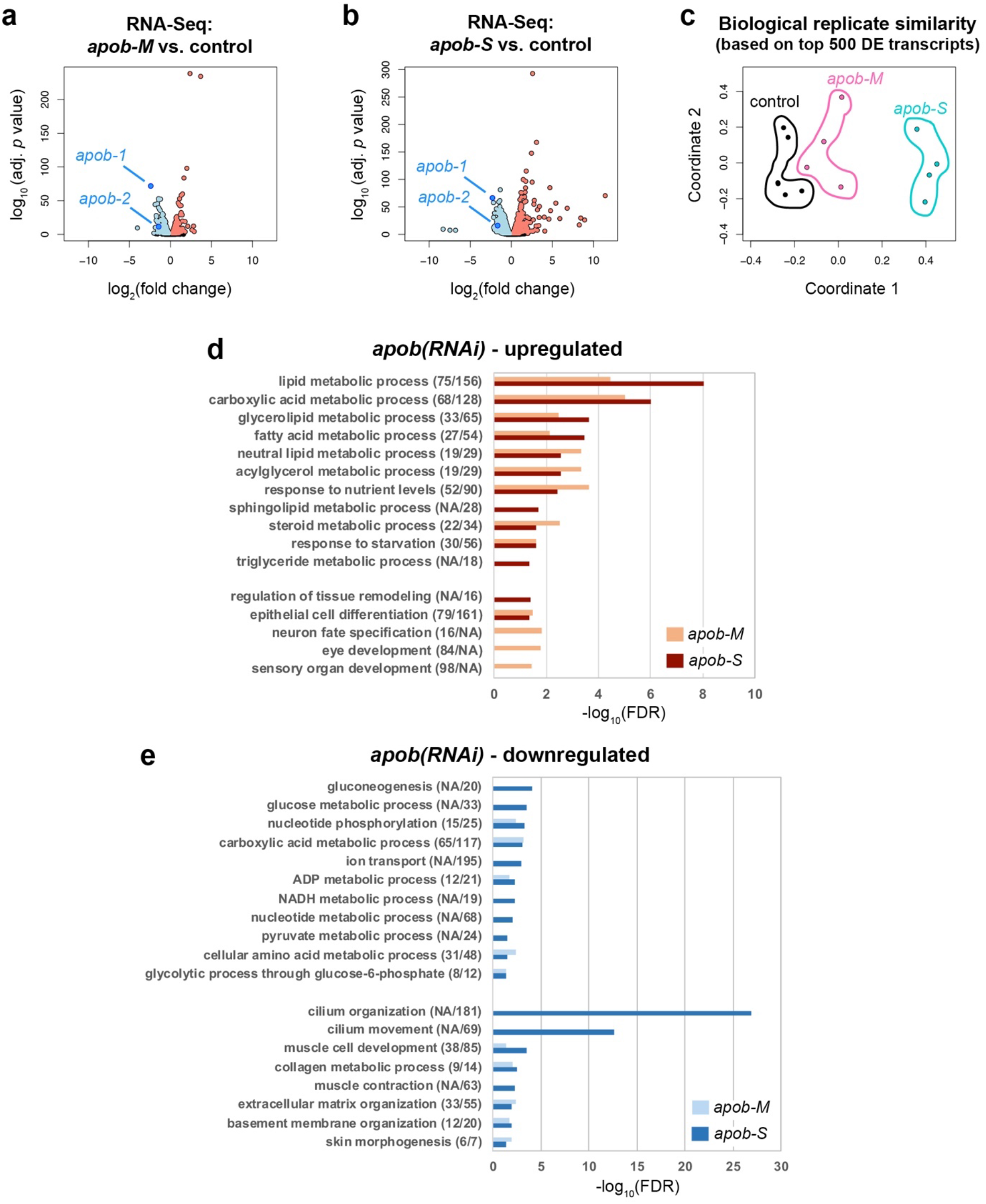
*apob* RNAi dysregulates transcripts involved in metabolism- and differentiation-related processes. **(a-b)** Volcano plots showing significantly downregulated (blue) and upregulated (pink) transcripts in *apob-M* **(a)** and *apob-S* **(b)** uninjured animals. **(c)** Multi-dimensional scaling plot showing similarity of control and RNAi sample libraries, using the biological coefficient of variation method to calculate distances between each library based on the 500 most variable transcripts across all samples. **(d-e)** Gene Ontology Biological Process categories over-represented among transcripts upregulated **(d)** or downregulated **(e)** by *apob* RNAi. Numbers of transcripts dysregulated indicated in parentheses (*apob-M/apob-S*). NA, not applicable (GO category not enriched in *apob-M* or *apob-S*). - log10(FDR) on X-axis.

**Supplementary Figure 6.**
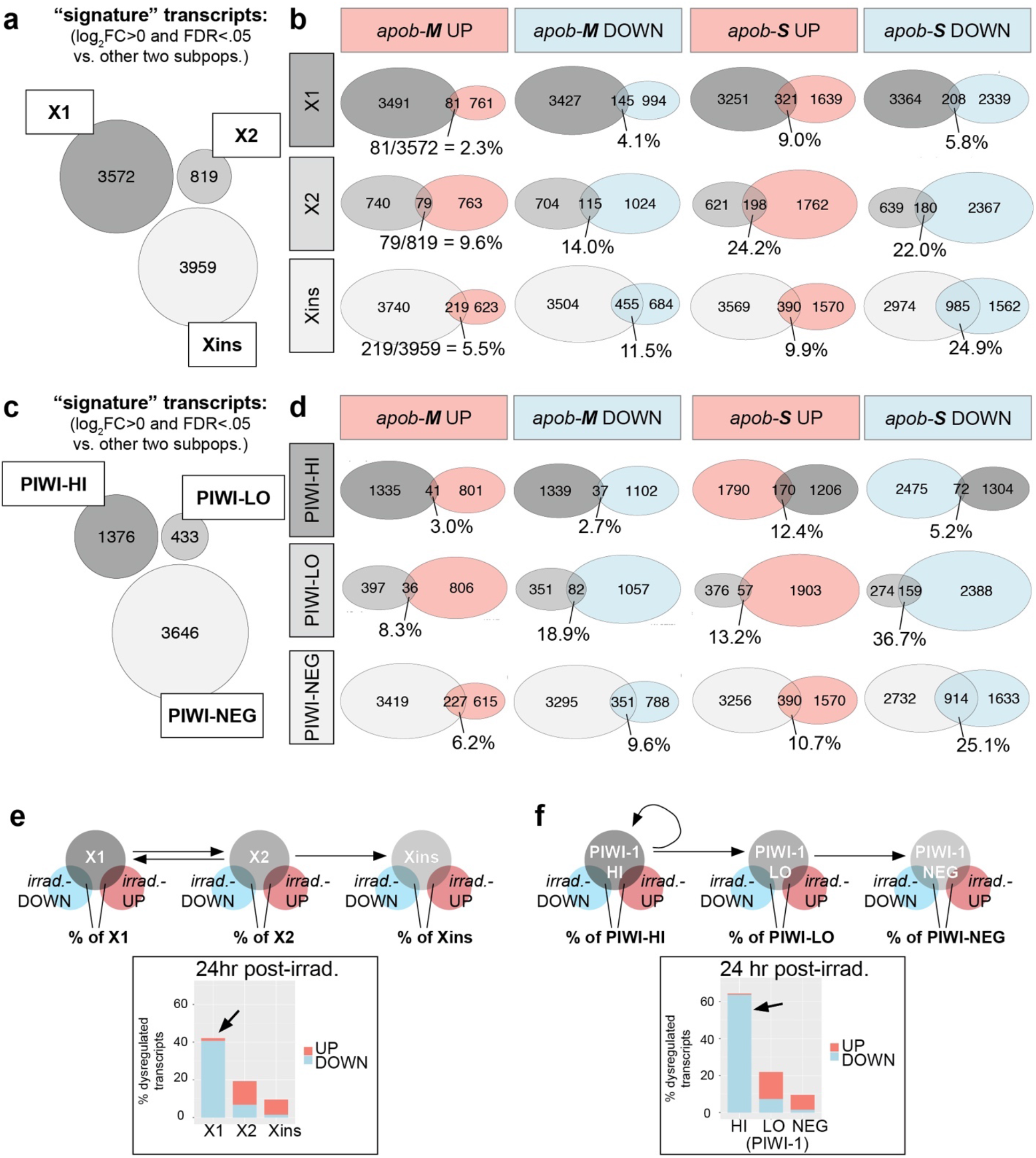
*apob* RNAi preferentially dysregulates transcripts in differentiating neoblast progeny and mature post-mitotic cells. **(a)** Venn diagram showing no overlap between X1, X2, and Xins “signature” transcripts (defined as having logFC>0 (FDR<.05) vs. transcripts in both other fractions). **(b)** Venn diagrams showing overlap between X1, X2, and Xins signature transcripts and transcripts up or down in *apob-M* and *apob-S*. Percentages of X1/X2/Xins dysregulated are indicated. **(c)** Venn diagram showing no overlap between PIWI-HI, PIWI-LO, and PIWI-NEG signature transcripts (defined as having logFC>0 (FDR<.05) vs. transcripts in both other fractions). **(d)** Venn diagrams showing overlap between PIWI-HI, PIWI-LO, and PIWI-NEG signature transcripts and transcripts up or down in *apob-M* and *apob-S*. Percentages of PIWI-HI/PIWI-LO/PIWI-NEG dysregulated are indicated. **(e-f)** X1-enriched **(e)** and PIWI-HI-enriched **(f)** signature transcripts were preferentially downregulated in planarians 24 hr post-irradiation. Venn diagrams at top show analysis schemes: percent of X1/X2/Xins **(e)** and PIWI-HI/PIWI-LO/PIWI-NEG **(f)** signature transcripts that overlap with transcripts dysregulated in planarians 24 hr post-irradiation. Histograms show percentage of signature transcripts up- and down-regulated in planarians 24 hr post-irradiation. Analysis of data from ^49^ (see Methods).

**Supplementary Figure 7.**
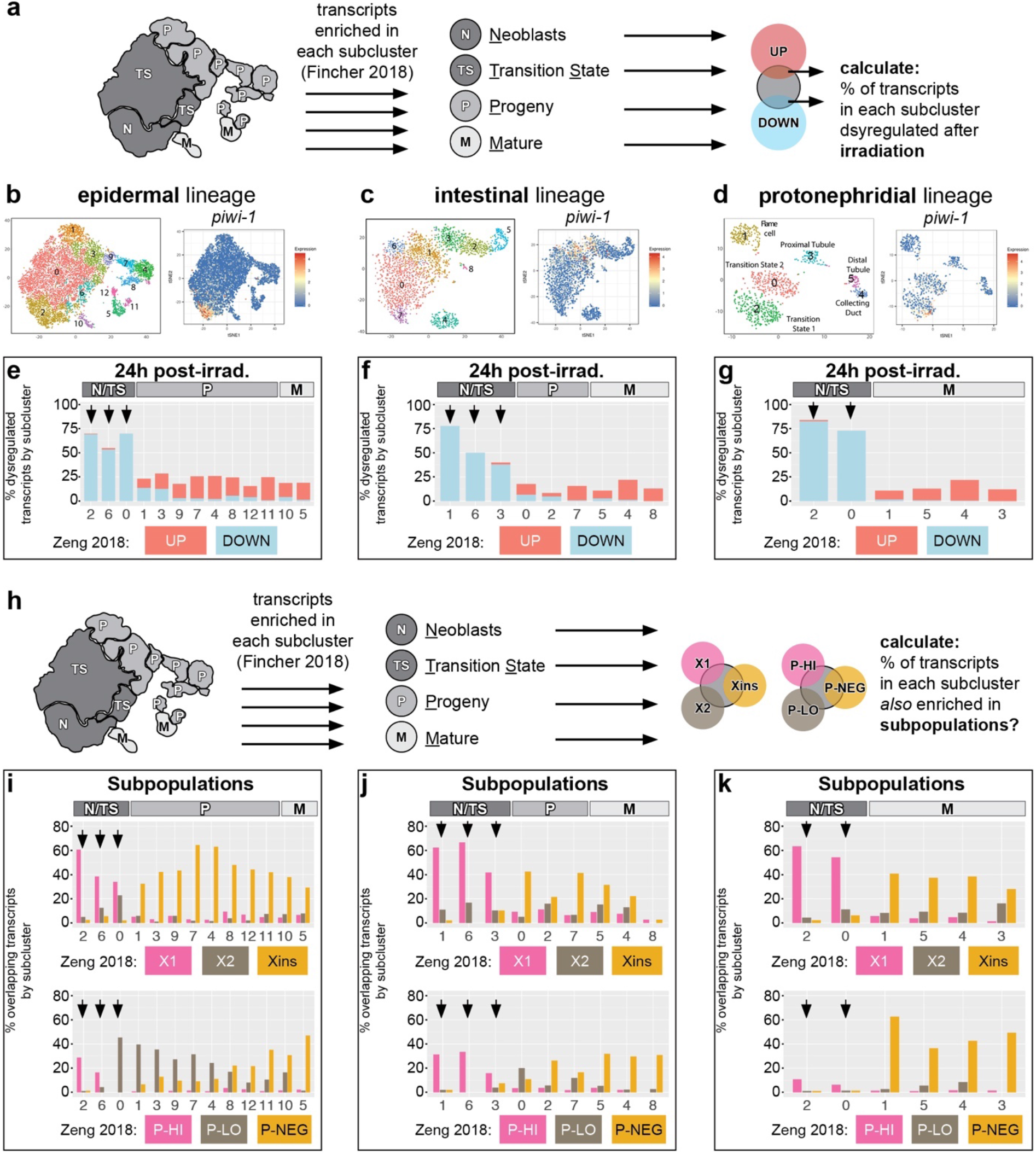
Mapping of transcripts from 24 hr post-irradiation and cell-state-enriched subpopulations to single-cell subclusters in three lineages. **(a)** Schematic example (for epidermal lineage) illustrating how transcripts dysregulated in 24 hr post-irradiation animals^49^ were cross-referenced with neoblast (N), transition state (TS), progeny (P), and mature (M) cell state subclusters^40^. See Methods for details. **(b-d)** t-SNE plots (digiworm.wi.mit.edu) indicate subclusters and *piwi-1* mRNA expression for each lineage. **(e-g)** Cross-referencing of scRNA-Seq data^40^ with RNA-Seq data from whole planarians 24 hr post-irradiation^49^. Histograms showing that irradiation preferentially caused downregulation of transcripts enriched in neoblast (”N“) and transition state (“TS”) subclusters in multiple cell type lineages (arrows), by contrast to the effects of *apob* RNAi (see also Fig. 5). **(h)** Cell state schematic and Venn diagrams show analysis strategy to calculate proportion of subcluster-enriched transcripts^40^ also enriched in sorted planarian cell subpopulations^49^. **(i-k)** Cross-referencing of scRNA-Seq data^40^ with bulk sorted cell RNA-Seq data^49^. Histograms showing that X1/PIWI-HI (“P-HI”) signature transcripts were primarily enriched in neoblast (“N”) and transition state (“TS”) subclusters, and that Xins/PIWI-NEG (“P-NEG”) signature transcripts were enriched in progeny (“P”) and mature (“M”) cell state subclusters. In epidermal and intestinal lineages, X2/PIWI-LOW (“P-LO”) signature transcripts were most highly enriched in neoblast/transition state and progeny subclusters. In the protonephridial lineage X2/PIWI-LO transcripts were more uniformly distributed, possibly due to fewer subclusters (and/or lower resolution of transition states) in this scRNA-Seq dataset.

**Supplementary Figure 8.**
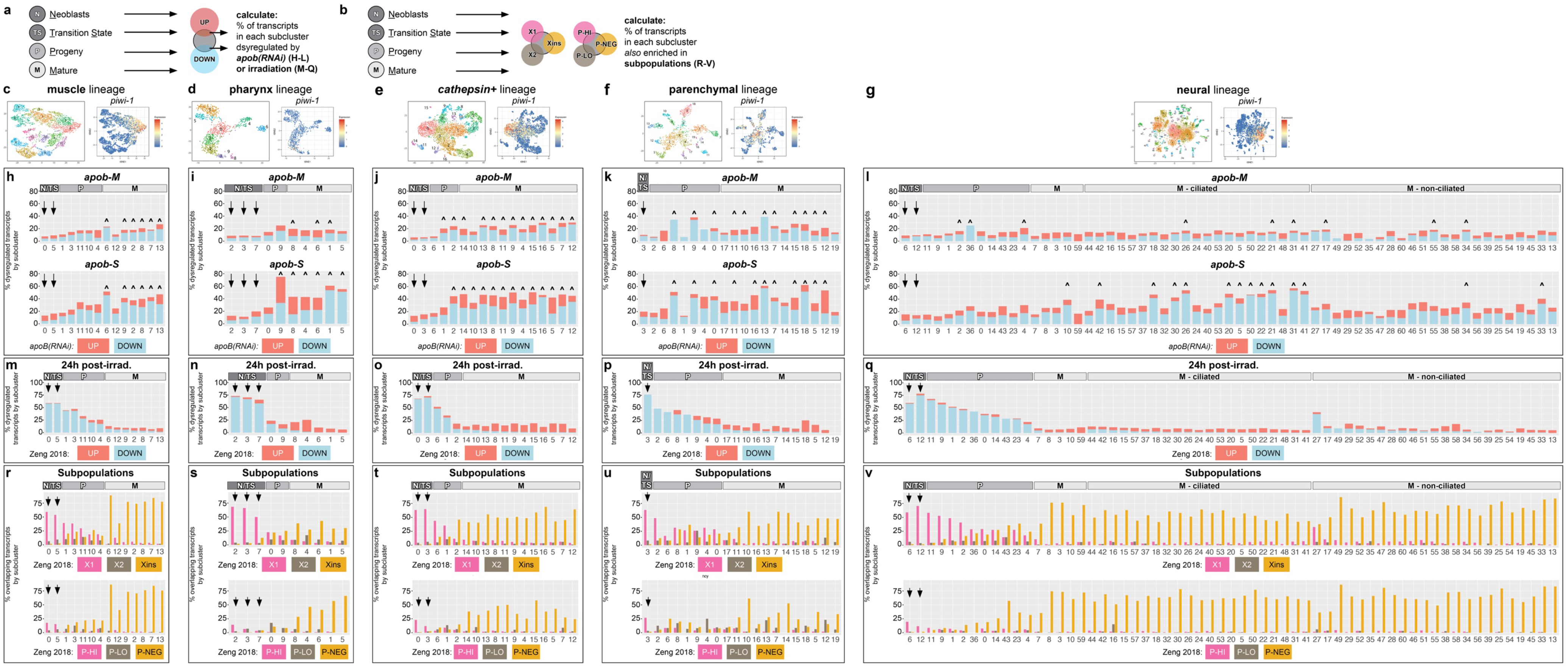
Additional examples of dysregulation of transcripts in differentiating neoblast progeny and mature cells by *apob* RNAi. **(a)** Generic scheme used to identify overlap of transcripts enriched in specific subclusters/cell states^40^ that were dysregulated in *apob(RNAi)* planarians (this study) or 24 hr post-irradiation^49^. **(b)** Generic scheme to calculate proportion of cell-state-enriched transcripts^40^ also enriched in sorted planarian cell subpopulations^49^. **(c-g)** t-SNE plots (digiworm.wi.mit.edu) indicate subclusters and *piwi-1* mRNA expression for each lineage. **(h-l)** *apob* knockdown dysregulated greater proportions of transcripts in progeny (“P”) and mature (“M”) subclusters in multiple cell type lineages. Arrows indicate less-affected transcripts in neoblast/transition state (“N/TS”) subclusters. **(m-q)** Cross-referencing of scRNA-Seq data^40^ with RNA-Seq data from whole planarians 24 hr post-irradiation^49^. Transcripts enriched in neoblasts/transition state subclusters were preferentially downregulated 24 hr post-irradiation (arrows), by contrast to the effects of *apob* RNAi (H-L). **(r-v)** Cross-referencing of scRNA-Seq data^40^ with bulk sorted cell RNA-Seq data^49^. Histograms showing that X1/PIWI-HI (“P-HI”) signature transcripts were primarily enriched in neoblast (“N”), transition state (“TS”), and progeny (“P) subclusters; that X2/PIWI-LOW (“P-LO”) signature transcripts were most highly enriched in progeny subclusters; and that Xins/PIWI-NEG (“P-NEG”) signature transcripts were enriched in mature (“M”) cell state subclusters. Carets (^) indicate significant gene expression overlap (**h-l**, *p<*0.05, Fisher’s exact test, see Source Data 2 for individual *p* values).

**Supplementary Figure 9.**
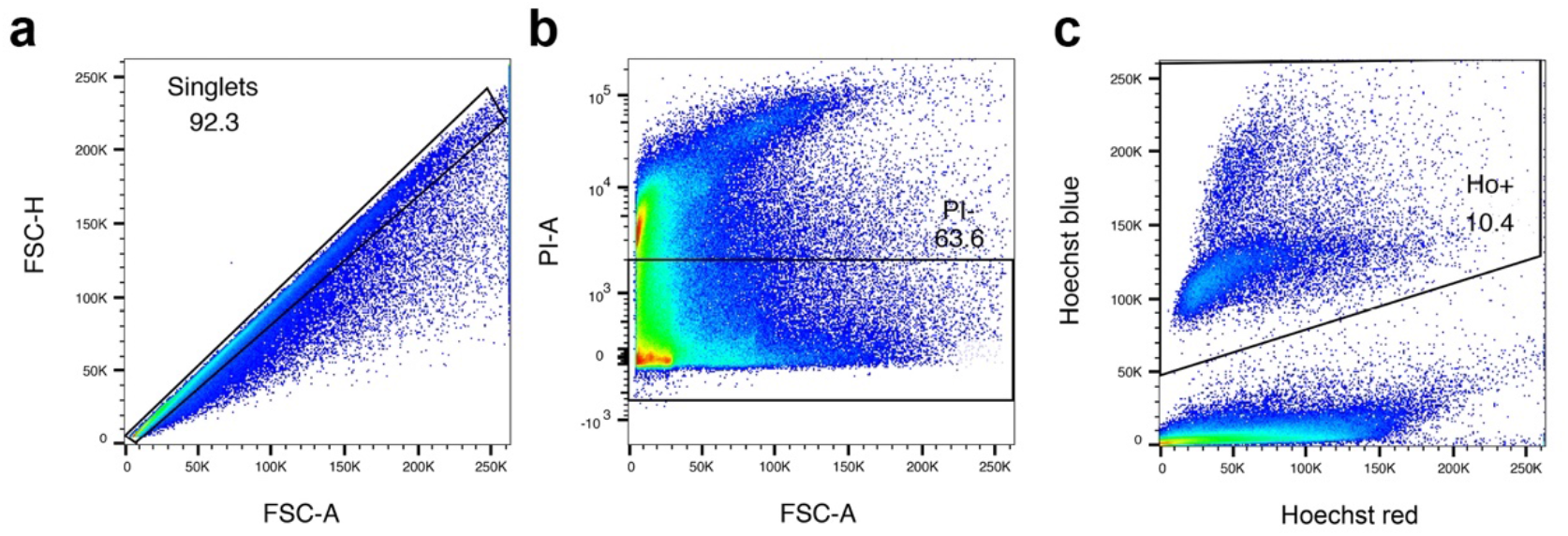
Gating strategy for flow cytometry experiments. **(a)** Forward scatter height (FSC-H) vs. forward scatter area (FSC-A) gate to limit to singlet events. **(b)** Propidium iodide (PI-A) vs. forward scatter area (FSC-A) gate to limit to PI negative (e.g., non-dead) events. **(c)** Hoechst 33342 blue (y-axis) vs. Hoechst 33342 red (x-axis) gates to limit to Hoechst-positive (e.g., ≥2C DNA content) events. Percentages of events after each gating step are indicated.

## Supplementary Data

**Supplementary Data 1. Up- and downregulated transcripts in *nkx2.2(RNAi)* planarians. (a)** RNA-Seq data for all transcripts detected in uninjured control vs. *nkx2.2(RNAi)* planarians, together with best BLAST hits for human/mouse/zebrafish/fly/*C. elegans*. **(b)** RNA-Seq data for 766 significantly downregulated transcripts in *nkx2.2(RNAi)* relative to control planarians. **(c)** RNA-Seq data for 719 significantly upregulated transcripts in *nkx2.2(RNAi)* relative to control planarians.

**Supplementary Data 2. Gene Ontology Biological Process terms enriched for transcripts up- and downregulated by *nkx2.2* RNAi. (a)** Biological Process term enrichment data for 766 transcripts downregulated in *nkx2.2(RNAi)* planarians. **(b)** Biological Process term enrichment data for 719 transcripts upregulated in *nkx2.2(RNAi)* planarians.

**Supplementary Data 3. Transcripts annotated with lipid-related Gene Ontology terms in whole fragment planarian regeneration transcriptome. (a)** Transcripts annotated with the term “lipid transport.” **(b)** Transcripts annotated with the term “triglyceride.” **(c)** Transcripts annotated with the term “cholesterol.” Expression data are derived from NCBI GEO GSE107874 ^49^ mapped to the dd_Smed_v6 transcriptome ^105^ (see Methods). Some transcripts may be annotated with multiple GO terms sharing the key words indicated, and therefore have multiple entries.

**Supplementary Data 4. Up- and downregulated transcripts in *apob(RNAi)* “mild” and “severe” planarians. (a)** RNA-Seq data for all transcripts detected in uninjured control vs. *apob-1(RNAi);apob-2(RNAi)* planarians, together with best BLAST hits for human/mouse/zebrafish/fly/*C. elegans*. **(b)** RNA-Seq data for 842 significantly upregulated transcripts in *apob-M(RNAi)* relative to control planarians. **(c)** RNA-Seq data for 1960 significantly upregulated transcripts in *apob-S(RNAi)* relative to control planarians. **(d)** RNA-Seq data for 1139 significantly downregulated transcripts in *apob-M(RNAi)* relative to control planarians. **(e)** RNA-Seq data for 2547 significantly downregulated transcripts in *apob-S(RNAi)* relative to control planarians.

Supplementary Data 5. Gene Ontology Biological Process terms enriched for transcripts up- and downregulated by *apob* RNAi. (a) Biological Process term enrichment data for 842 transcripts upregulated in *apob-M(RNAi)* planarians. **(b)** Biological Process term enrichment data for 1960 transcripts upregulated in *apob-S(RNAi)* planarians. **(c)** Biological Process term enrichment data for 1139 transcripts downregulated in *apob-M(RNAi)* planarians. **(d)** Biological Process term enrichment data for 2547 transcripts downregulated in *apob-S(RNAi)* planarians.

**Supplementary Data 6. Gene and transcript identities used in phylogenetic analyses and gene expression studies. (a)** ApoB and related homologs used for domain diagram and phylogenetic analysis. **(b)** Alignment of N-terminal Vitellogenin domains used for phylogenetic tree. **(c)** Lipoprotein receptor homologs used for domain diagram. **(d)** Transcript identities, sequences, cloning primers and vectors used in this study.

**Source Data 1. Differential expression analysis of neoblast subpopulations from GSE107874.**

**Source Data 2. Numerical data and statistical analyses.**

## Notes

### Competing Interest Statement

The authors have declared no competing interest.

